# The angiosperm seed life cycle follows a developmental reverse hourglass

**DOI:** 10.1101/2024.12.20.629609

**Authors:** Asif Ahmed Sami, Leónie Bentsink, Mariana A. S. Artur

## Abstract

The seed life cycle is one of the most crucial stages in determining the ecological success of angiosperms. It broadly comprises three developmental phases - embryogenesis, maturation, and germination. Among these phases, seed maturation is particularly critical, serving as a bridge between embryo development and germination. During this phase, seeds accumulate nutrient reserves and acquire essential physiological traits, such as desiccation tolerance, vital for seed survival in diverse environments. Phylotranscriptomics in *Arabidopsis thaliana* has shown that embryogenesis and germination follow an hourglass-like development, with high expression of older and conserved genes at the mid-developmental stages. However, unlike embryogenesis and germination, a phylotranscriptomic study of seed maturation has not yet been performed and a comprehensive overview of the phylotranscriptomic landscape throughout the entire seed life cycle is still lacking. Here, we combined existing RNA-seq data covering all three phases of the *Arabidopsis* seed life cycle to construct a complete picture of the phylotranscriptomic pattern of the seed life cycle by generating transcriptome age index (TAI) and transcriptome divergence index (TDI) profiles. We found that the seed life cycle resembles a reverse hourglass-like pattern, with seed maturation exhibiting increased expression of younger genes with divergent expression patterns compared to embryogenesis and germination. Notably, this pattern of increased expression of younger genes during seed maturation is also conserved across both dicot and monocot species. Tissue-specific phylotranscriptomic analyses revealed that, in monocots, the increased expression of younger genes during maturation is largely driven by genes expressed in the endosperm. Overall, our findings highlight the major shifts in phylotranscriptomic patterns during the seed life cycle and establish seed maturation as a pivotal developmental phase enabling the expression of young and rapidly evolving genes critical for seeds’ adaptive capacity in their surrounding environment.

## Introduction

The evolution of the seed habit is a major innovation in the plant kingdom and is regarded as one of the most successful means of sexual reproduction in plants (Haig and Westoby, 1989). Seeds have facilitated the large-scale diversification of plants and paved the way for gymnosperms and angiosperms to dominate global ecosystems (Linkies et al., 2010). The seed life cycle can be broadly divided into three phases: embryogenesis, maturation, and germination (Kermode, 1990; Holdsworth et al., 1999). Embryogenesis marks a series of coordinated cell divisions that establish the basic body plan of the plant. Subsequently, seeds transition into the maturation phase, a highly physiologically active phase whereby seeds accumulate nutrient reserves (e.g. oils, sugars, and proteins) and acquire quality traits such as germinability, dormancy, desiccation tolerance, and longevity which are vital for their survival in diverse environments (Kermode, 1990). Post maturation, most seeds enter a quiescent dry state whereby they can persist in the environment for prolonged periods before conditions are favourable for germination to commence.

On the contrary, in most animals, embryonic development is a continuous process devoid of a quiescent maturation phase. Animal embryogenesis follows an hourglass pattern of development, whereby embryos of different species exhibit morphological divergence during early and later stages of development but converge towards higher resemblance during mid-embryonic development (referred to as the phylotypic stage) (Galis and Metz, 2001; Willmore, 2012; Irie and Kuratani, 2014; Drost et al., 2017). Several factors lead to evolutionary constraints that converge in higher morphological similarity during mid-embryogenesis compared to earlier or later developmental stages. Evidence of the embryonic hourglass was also substantiated on a transcriptome level using phylotranscriptomic indices (Domazet-Lošo and Tautz, 2010; Quint et al., 2012). These indices combine the evolutionary distance of genes with their expression levels to determine the transcriptome age during a specific stage of development. Two commonly used indices include the transcriptome age index (TAI) and transcriptome divergence index (TDI), which rely on evolutionary age and sequence divergence, respectively. TAI and TDI patterns were used to demonstrate the presence of a molecular hourglass signature in animals, plants, fungi, and brown algae using both bulk and single-cell RNA-seq (Domazet-Lošo and Tautz, 2010; Quint et al., 2012; Cheng et al., 2015; Ma and Zheng, 2023; Wu et al., 2024; Lotharukpong et al., 2024). Among angiosperms, the embryonic hourglass was shown for multiple species, including *Arabidopsis thaliana* L. (hereby referred to as *Arabidopsis*) (Quint et al., 2012), *Brassica* spp. (Gao et al., 2022), wheat (Xiang et al., 2019), and *Zea mays* L. (Wu et al., 2024). Intriguingly, the hourglass pattern was also reported post-embryonically, including seed germination of *Arabidopsis* (Drost et al., 2016). The existence of dual hourglass-like patterns during the seed life cycle suggests the presence of evolutionary forces that drive the expression of conserved genes in both phases.

Unlike embryogenesis and germination, a phylotranscriptomic study of seed maturation has not yet been conducted, likely due to the previous lack of high-resolution temporal RNA-seq data that captures the transcriptional landscape of this phase in *Arabidopsis* (Artur et al., 2024). Consequently, a complete picture of the phylotranscriptomic pattern of the entire seed life cycle is still missing. Seed maturation is a particularly critical phase of the seed life cycle, as it serves as a bridge between embryo development and germination and where the developmental fitness, plasticity, and ecological success of seeds are mostly determined. Thus, understanding the dynamics and drivers of evolution during phase transitions of the entire seed life cycle can shed light on the ecological importance of physiological adaptations in seeds.

In this work, we examined the phylotranscriptome landscape of *Arabidopsis* seed life cycle to identify the evolutionary patterns across key phase transitions. We found that, unlike embryogenesis and germination, the *Arabidopsis* maturation transcriptome is characterized by high TAI and TDI values, with the phylotranscriptomic pattern of the entire seed life cycle resembling a reverse hourglass. Leveraging RNA-seq datasets from multiple studies, we generated the first phylotranscriptomic profile of the seed life cycle across multiple angiosperm species, including monocots and dicots. Remarkably, we identified a conserved high TAI pattern during maturation in both monocot and dicot lineages, with the endosperm significantly contributing to the high TAI during maturation of monocots. Comparison of seed maturation with pollen development, another phase with high TAI and a reverse hourglass pattern (Wu et al., 2014; Cui et al., 2015; Gossmann et al., 2016a; Julca et al., 2021), revealed that the younger genes driving the reverse hourglass pattern are unique to the seed transcriptome. Overall, our findings establish seed maturation as a pivotal developmental phase enabling the expression of young genes and rapidly evolving genes critical for seeds’ adaptive capacity in its surrounding environment.

## Results

### Transcriptome age and divergence indices reveal a reverse hourglass pattern during the *Arabidopsis* seed life cycle

The developmental hourglass pattern during embryogenesis and germination is well-documented in *Arabidopsis* (Quint et al., 2012; Drost et al., 2016). However, to identify the evolutionary patterns across key phase transitions of the entire seed life cycle, a high-resolution temporal seed maturation transcriptome was essential. We therefore combined recently available RNA-seq data on *Arabidopsis* seed maturation (Artur et al., 2024) with previously generated RNA-seq data of embryogenesis (Hofmann et al., 2019) and germination. Together, these three RNA-seq datasets cover the complete life cycle of *Arabidopsis* seeds, from the preglobular embryonic stage to 72 hours of germination (Supplemental Table 1). By executing homology searches against the proteomes of 4,940 species (including bacteria, archaea, fungi, animals, and plants) we classified *Arabidopsis* genes into different phylostrata (PS) based on their originating nodes (Supplemental Table 2). This resulted in a phylostratiographic map that grouped all *Arabidopsis* genes (27,416 genes) into discrete age groups ranging from PS1 to PS16 (Fig. 1a and Supplemental Table 3), with PS1 representing the oldest and PS16 the youngest phylostrata. The transcriptome age index (TAI) measure relies on the phylogenetic rank of a gene and its expression at a given developmental stage (Domazet-Lošo and Tautz, 2010) and was calculated for each sample in the three datasets (Supplemental Table 1). We visualized the TAI profiles for each phase individually and observed that both embryogenesis and germination displayed a significant hourglass pattern (*p_hg_* = 0.000116 and 0.0159, respectively), with a characteristic phylotypic phase during mid-development as previously described (Supplemental Fig. 1a,c) (Quint et al., 2012; Drost et al., 2016). In contrast, the TAI profile of the maturation phase did not deviate significantly from a flat line, indicating little variation in transcriptome age and suggesting a stable transcriptome profile throughout this phase (Supplemental Fig. 1b, *p_flt_* = 0.776). Plotting the TAI values of the maturation stages alongside those of embryogenesis and germination revealed a strikingly high TAI during seed maturation, distinguishing it from its adjacent phases (Fig. 1b). Intriguingly, the TAI profile of the entire seed life cycle significantly (*p_rev_* = 9.89 X 10^-5^) resembled a reverse hourglass pattern when maturation was considered the mid-developmental phase (Fig. 1b).

**Fig. 1.**
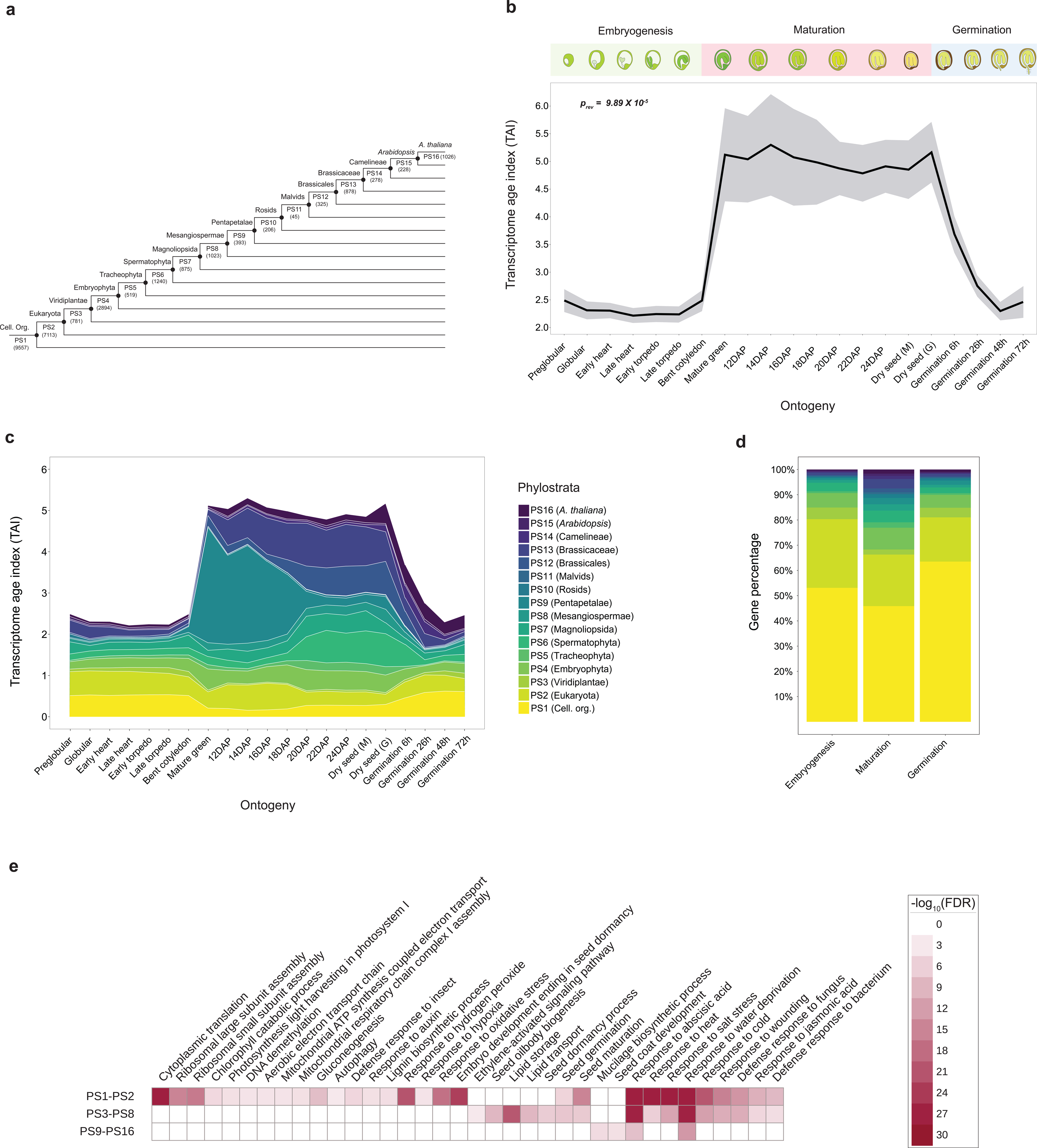
TAI profile throughout the seed life cycle of *Arabidopsis*. **a,** Phylostratigraphy of *Arabidopsis* showing the distribution of protein-coding genes over sixteen phylostrata (PS1-PS16). **b**, TAI pattern during three distinct phases (embryogenesis, maturation, and germination) of the *Arabidopsis* seed life cycle. Colored boxes correspond to stages that belong to each phase. Shaded regions around the TAI line indicate the standard deviation (s.d.) calculated from 1000 permutations. Embryogenesis, maturation, and germination stages were defined as early, mid, and late developmental stages, respectively, during the reverse hourglass test. The p-value indicates the significance of the reverse hourglass test. DAP stands for days after pollination. **c**, Contribution of individual phylostratum to the overall TAI profile. Phylostratum are arranged from youngest to oldest from top to bottom. The TAI values are not cumulative, and each band indicates the contribution of a single phylostratum alone. **d**, Percentage of genes from each phylostratum among the top 5% most expressed genes for each phase. For each phase, genes were filtered on a threshold of average TPM ≥ 1 and then sorted from highest to lowest expression to select the top 5% of genes. **e**, GO terms enriched among the top 5% maturation genes. Genes were grouped into three groups - PS1-PS2, PS3-PS8, and PS9-PS16.

While TAI captures ancestral distance-based evolutionary signals, the transcriptome divergence index (TDI) depicts evolutionary signals from recent sequence divergence (Quint et al., 2012; Drost et al., 2015). We then calculated the TDI per RNA-seq sample by combining Ka/Ks (non-synonymous / synonymous substitutions) ratio of each gene with its expression level (Supplemental Table 4). Consistent with the TAI profile, the TDI profile also resembled a reverse hourglass (*p_rev_* = 0.00783, Supplemental Fig. 1d), with maturation exhibiting higher TDI values than the adjacent phases. Taken together, the TAI and TDI profiles suggest that embryogenesis and germination exhibit relatively lower TAI and TDI values, likely due to greater reliance on more conserved or less rapidly evolving genes (Quint et al., 2012; Drost et al., 2016). On the other hand, the seed maturation transcriptome of *Arabidopsis* exhibits the opposite trend, characterized by the contribution of younger and more rapidly evolving genes.

### Genes underlying the high maturation TAI are associated with seed maturation traits

To uncover the underlying drivers of the reverse hourglass pattern of the seed life cycle we explored the TAI contributions of individual phylostrata. The two oldest phylostrata, i.e., PS1 and PS2, contributed the highest with the TAI during embryogenesis and germination (Fig. 1c and Supplemental Fig. 2). However, there was a noticeable decline in their contribution during the transition from embryogenesis (bent cotyledon stage) to maturation (mature green stage) (Supplemental Fig. 3a). In contrast, the contribution of several younger phylostrata, such as PS6 (Spermatophyta), PS9 (Pentapetalae), PS12 (Brassicales), and PS13 (Brassicaceae), was higher during maturation in relation to embryogenesis (Fig. 1c). Upon examining the relative expression levels of all phylostrata, we observed a consistent pattern: older phylostrata exhibited higher relative expression during embryogenesis and germination, while younger phylostrata had increased relative expression primarily during maturation (Supplemental Fig. 3a-c). For instance, the contribution of PS9 (Pentapetalae) showed a steep increase at the onset of maturation, with a high TAI contribution starting from the bent cotyledon stage until 18 days after pollination (DAP) (Fig. 1c and Supplemental Fig. 2). By sorting the genes based on their average expression during the maturation (mature green to dry seeds), we discovered that some of the most highly expressed genes from PS9 were several known seed storage proteins such as seed storage albumins (1-4) and late embryogenesis abundant (LEA) proteins (AT2G21490 - dehydrin LEA, AT3G50980 - *XERO1*, and AT2G42560 - *LEA25*) (Supplemental Table 5). The high expression of seed storage protein-encoding genes from PS9 clearly coincides with the initiation of seed-filling during maturation (Baud et al., 2002). Likewise, the contribution of PS12 (Brassicales) and PS13 (Brassicaceae) also showed an increase during early maturation but reached the highest magnitude at later stages of maturation (18 DAP onwards) (Fig. 1c). For these two PS, the most highly expressed genes also included LEAs and dehydrins, along with several glycine-rich and proline-rich proteins, as well as hydroxyproline-rich glycoprotein family proteins (Supplemental Table 5). The contributions of PS6 (Spermatophyta) and PS7 (Magnoliopsida) during maturation were also noticeable, particularly at later maturation time points. The highest expressed genes from these two PS included defensins, LEAs, and lipid transport proteins (see Supplemental Table 5), which are well-known for their role in seed filling and desiccation tolerance (Angelovici et al., 2010; Verdier et al., 2013; Leprince et al., 2016).

This insight from individual phylostrata contributions prompted us to delve further into the specific genes and pathways shaping the seed maturation phylotranscriptome landscape. We hypothesized that the high expression of younger PS genes during maturation drives the reverse hourglass pattern in the seed life cycle. To test this, we first recalculated the TAI of the whole seed life cycle by removing either the top 1% (160 genes), 2% (320 genes), 5% (801 genes), 10% (1602 genes), or 20% (3203 genes) genes with the highest expression (average transcripts per million, TPM) during maturation only (Supplemental Table 6). Then, we tested whether removing these sets of genes affects the significance of the reverse hourglass pattern (Supplemental Fig. 4). We found that removing the top 5% genes was sufficient to lead to a non-significant reverse hourglass pattern (*p_rev_* = 0.0724), indicating that these genes are primarily responsible for the reverse hourglass pattern during the seed life cycle. Comparing the top 5% highly expressed genes in maturation with those in embryogenesis and germination revealed that the maturation phase had the highest percentage of younger PS genes, although a substantial portion of these genes still belonged to PS1 and PS2 (Fig. 1d). Interestingly, the expression (mean log10 TPM) of these top 5% maturation genes, especially from younger PS, was relatively more variable and in some cases higher than that of the top 5% genes of embryogenesis and germination (Supplemental Fig. 5). These findings suggest that seed maturation contains a higher proportion of younger PS genes with higher expression variation than embryogenesis and germination, highlighting their key role in driving the reverse hourglass of the seed life cycle.

We next performed gene ontology (GO) enrichment analysis to identify the biological processes that are associated with the top 5% maturation-expressed genes from the different PS. For this, we first classified all phylostrata into three groups – i) ‘older’ (PS1 and PS2, genes conserved among cellular organisms and eukaryotes), ii) ‘intermediate’ (PS3 to PS8, genes that originated with the green lineage until the origin of core angiosperms), and iii) ‘younger’ (PS9 to PS16, all genes that originated afterward). As expected, genes from older PS (PS1 and PS2) were specifically highly enriched for processes related to basic cellular functions such as translation, respiration, autophagy, and response to oxidative stress, as well as some specialized functions such as photosynthesis, defense, and lignin biosynthesis (Fig. 1e). We also found a high enrichment of genes from the older PS group involved in embryo development ending with seed dormancy, suggesting an ancestral origin of embryo development genes that are highly expressed during maturation. Looking at the number of GO annotation terms assigned to genes from different PS, we found that the older PS group have significantly higher number of annotated genes compared to the intermediate (P3-P8) and younger (P9-P16) PS groups (Supplemental Fig. 6). Despite this GO annotation bias, we found that seed maturation-related processes such as oil body biogenesis, lipid storage and transport and seed dormancy were exclusively enriched in genes from the intermediate PS group (PS3-PS8) and processes associated with mucilage biosynthesis and seed coat development were exclusively enriched in genes from the younger PS group (PS9-PS16) (Fig. 1e). We also found that two seed-specific processes, seed germination and seed maturation, were enriched in both older and intermediate PS. Strikingly, we found that two processes of notable importance for seed maturation and desiccation, response to abscisic acid (ABA) and response to water deprivation, were highly enriched in genes from all PS groups. Together, our findings indicate that seed maturation-enriched biological processes have multiple origins.

### The reverse hourglass of seed maturation is conserved across multiple plant species

To examine whether high TAI values during seed maturation and the reverse hourglass pattern of the seed life cycle observed for *Arabidopsis* is conserved across angiosperms, we investigated the transcriptome of phases of the seed life cycle of three crop species: the dicots brassica *(Brassica napus,* Brassicacea) and tomato (*Solanum lycopersicum*, Solanaceae), and the monocot maize (*Zea mays*, Poaceae). For *B. napus,* we combined RNA-seq data covering embryogenesis, maturation, and germination (Supplemental Table 1) (Boter et al., 2019; Bianchetti et al., 2021; Gao et al., 2022). Using phylostrata information (Supplemental Fig. 7-8 and Supplemental Table 7) and gene expression, we constructed the TAI profile of *B. napus* seed life cycle. Overall, the pattern resembled that of *Arabidopsis*, with maturation time points (mature green until thermal time interval 9, see methods) exhibiting higher TAI compared to time points of the flanking phases (Fig. 2a). Testing the overall TAI profile for a reverse hourglass pattern yielded a significant p-value (*p_rev_* = 1.47 X 10^-5^), indicating that the *B. napus* seed life cycle also follows a reverse hourglass-like development. Delving further into the contribution of each phylostrata, we found that most of the phylostrata with a high TAI contribution during *Arabidopsis* seed maturation also had higher TAI values in *B. napus*. For instance, PS6, PS9, PS12, and PS13 all had a higher TAI contribution during part or in all the maturation time points in both species (Fig. 2b and Supplemental Fig. 8).

**Fig. 2.**
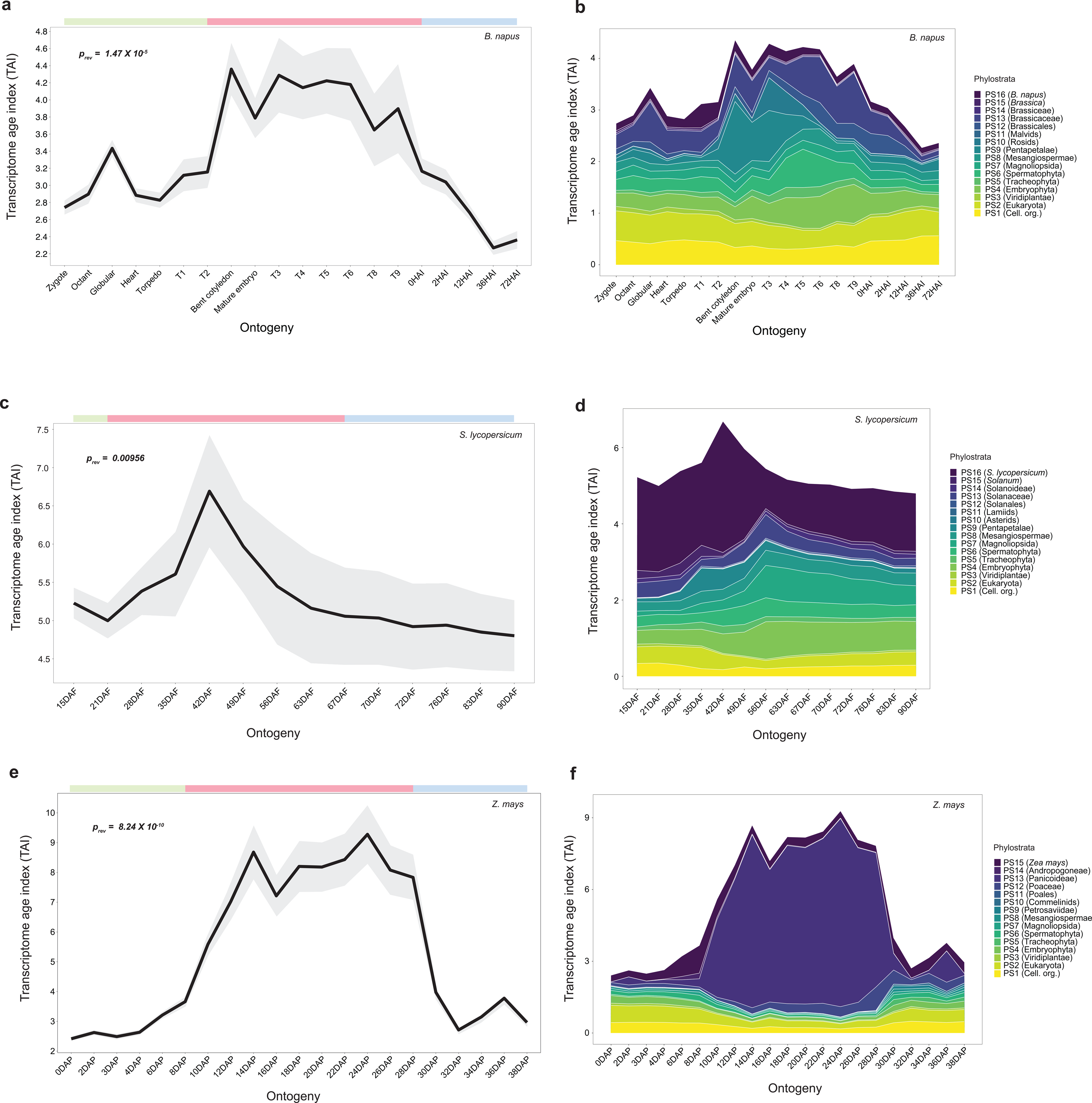
TAI profile during phases of the seed life cycle in three angiosperm species. **a**, TAI profile during the entire seed life cycle in *B. napus*. Green, red, and blue colored panel on top indicate stages that were considered part of early, mid, and late development, respectively, while performing the reverse hourglass test. The p-value indicates the significance of the test. T1-T9 (thermal time interval) indicates when seeds were harvested during maturation (Bianchetti et al., 2021). HAI indicates hours after imbibition. **b**, Contribution of individual phylostratum to the overall TAI profile in *B. napus*. **c**, TAI profile during parts of the seed life cycle in *S. lycopersicum*. Earlier stages (15 & 21 days after fertilization (DAF)) represent end of embryogenesis and 28 DAF marks initiation of seed filling, i.e., maturation (Bizouerne et al., 2021). **d**, Contribution of individual phylostratum to the overall TAI profile in *S. lycopersicum*. **e**, TAI profile during parts of the seed life cycle in *Z. mays*. **f**, Contribution of individual phylostratum to the overall TAI profile in *Z. mays*.

For tomato, we generated the phylostratigraphy (Supplemental Fig. 9 and Supplemental Table 7) and calculated the TAI values using RNA-seq data that covered end of embryogenesis and maturation phases (Bizouerne et al., 2021). We plotted the calculated TAI values over the different time points (days after flowering, DAF) (Fig. 2c). Despite the absence of early embryogenesis and germination time points, the overall TAI profile also followed a significant reverse hourglass pattern (*p_rev_* = 0.00956). Similar to our findings in Arabidopsis, analysis of individual phylostrata contributions revealed a high TAI from younger phylostrata during tomato maturation (28 DAF onwards) (Fig. 2d and Supplemental Figs. 10a-c). A striking feature of the tomato TAI profile was the high contribution of the youngest phylostrata (PS16) during the entire maturation phase. However, removing PS16 genes from the reverse hourglass test still resulted in a significant pattern (*p_rev_* = 0.00506) (Supplemental Fig. 11), indicating that high expression of PS16 genes is not the sole driver of the reverse hourglass pattern in tomato. Amongst the other phylostrata highly contributing to the tomato TAI pattern, PS4, PS6, PS7, and PS9 exhibited high TAI contributions either during early (28 DAF to 42 DAF) or late maturation (49 DAF onwards) (Fig. 2d and Supplemental Figs. 10a-b).

Considering our finding of a conserved reverse hourglass pattern in three dicots (*Arabidopsis*, *B. napus*, and tomato) we asked if a similar trend is also found in a monocot species. To test this, we calculated the TAI profile of maize using phylostrata information (Supplemental Fig. 12 and Supplemental Table 7) and existing RNA-seq datasets covering embryogenesis (0 DAP to 10 DAP) and maturation (12 DAP to 38 DAP) (Chen et al., 2014) (Supplemental Table 1). We found with high statistical significance that the reverse hourglass pattern was also observed when analysing maize seed embryogenesis and maturation phases together (Fig. 2e, *p_rev_* = 8.24 X 10^-10^). Strikingly, delving further into the contribution of individual phylostratum showed that the high TAI profile during maize seed maturation (12 DAP until 38 DAP) was largely due to a single phylostratum, PS13 (Panicoideae) (Fig. 2f and Supplemental Fig. 13c). We investigated the causal genes underlying this peak by looking at the average expression (TPM) during the same maturation time points. Interestingly, we identified several zein-encoding genes as the most highly expressed genes from PS13 (Supplemental Table 8 and 9). Zeins are widely known to be the most abundant storage proteins in maize seeds (Schmitz et al., 1997; Woo et al., 2001; Flint-Garcia et al., 2009). Although the removal of zein genes from PS13 did not affect the significance of the reverse hourglass pattern (*p_rev_* = 0.0416), it profoundly reduced the TAI peak during maturation (Supplemental Fig. 14a-b). In contrast, all other phylostratum ranging from PS1 to PS11 showed lower and an overall decline in their TAI values and relative expression during maturation (Supplemental Figs. 13a-b), whereas PS12, PS14, and PS15 showed a variable trend during the entire time course (Supplemental Fig. 13c). This was a clear deviation from the overall pattern seen for the three dicot species mentioned above, where younger phylostrata showed higher TAI values throughout seed maturation. Similar to the TAI profile, the TDI profile significantly followed a significant reverse hourglass pattern in *B. napus* and *Z. mays* (*p_rev_* = 0.0.0472 and *p_rev_* = 0.00352, respectively) (Supplemental Fig. 15a and c). However, this was not the case for *S. lycopersicum* (*p_rev_* = 0.937), likely due to the lack of a more complete embryogenesis time series (Supplemental Fig. 15b).

To assess the (dis)similarity of the seed maturation transcriptome between these different crop species and *Arabidopsis*, we determined the respective orthogroups of the genes from the four species and selected the top 5% genes which had the highest average expression values during maturation from each species (Supplemental Table 6 and 9). The expression of these top genes was averaged per orthogroup for every RNA-seq sample to attain expression for each orthogroup. This led to 681 orthogroups for *Arabidopsis*, 1020 for *B. napus*, 822 for *S. lycopersicum*, and 724 for *Z. mays*, with 124 orthogroups shared across the four species (Fig. 3a). A principal component analysis (PCA) based on the expression of these top orthogroups showed a separation of the *S. lycopersicum* samples in relation to the other species ‘samples (PC1 = 51.3% of variance), while the maturation time points cluster mostly according to the respective species (PC2 = 30.9% of variance) with very little overlap between maturation time points of different species (Fig. 3b). This shows that the maturation transcriptome is highly diverse when considering the top orthogroups. However, this separation largely disappears when the PCA was performed with only the 124 shared orthogroups (Supplemental Fig. 16). The orthogroups partially shared amongst the four species (referred to as non-core orthogroups) included genes from nearly all phylostrata in each species, except for PS11 genes in *B. napus* (Malvids) and of *Z. mays* (Poales) (Fig. 3c, left). The core maturation orthogroups (shared by all four species) comprised genes from only six distinct, mostly older, phylostrata. As expected, most of these genes originated from PS1 and PS2 (Fig. 3c, right) and were enriched for basic cellular processes (using *Arabidopsis* gene functions annotation), while genes from later phylostrata (PS4 to PS7) were enriched for seed-related functions (Supplemental Fig. 17a). On the other hand, genes from the non-core orthogroups were enriched for a range of diverse functions including primary metabolism, responses to stress, and seed development (Supplemental Fig. 17b).

**Fig. 3.**
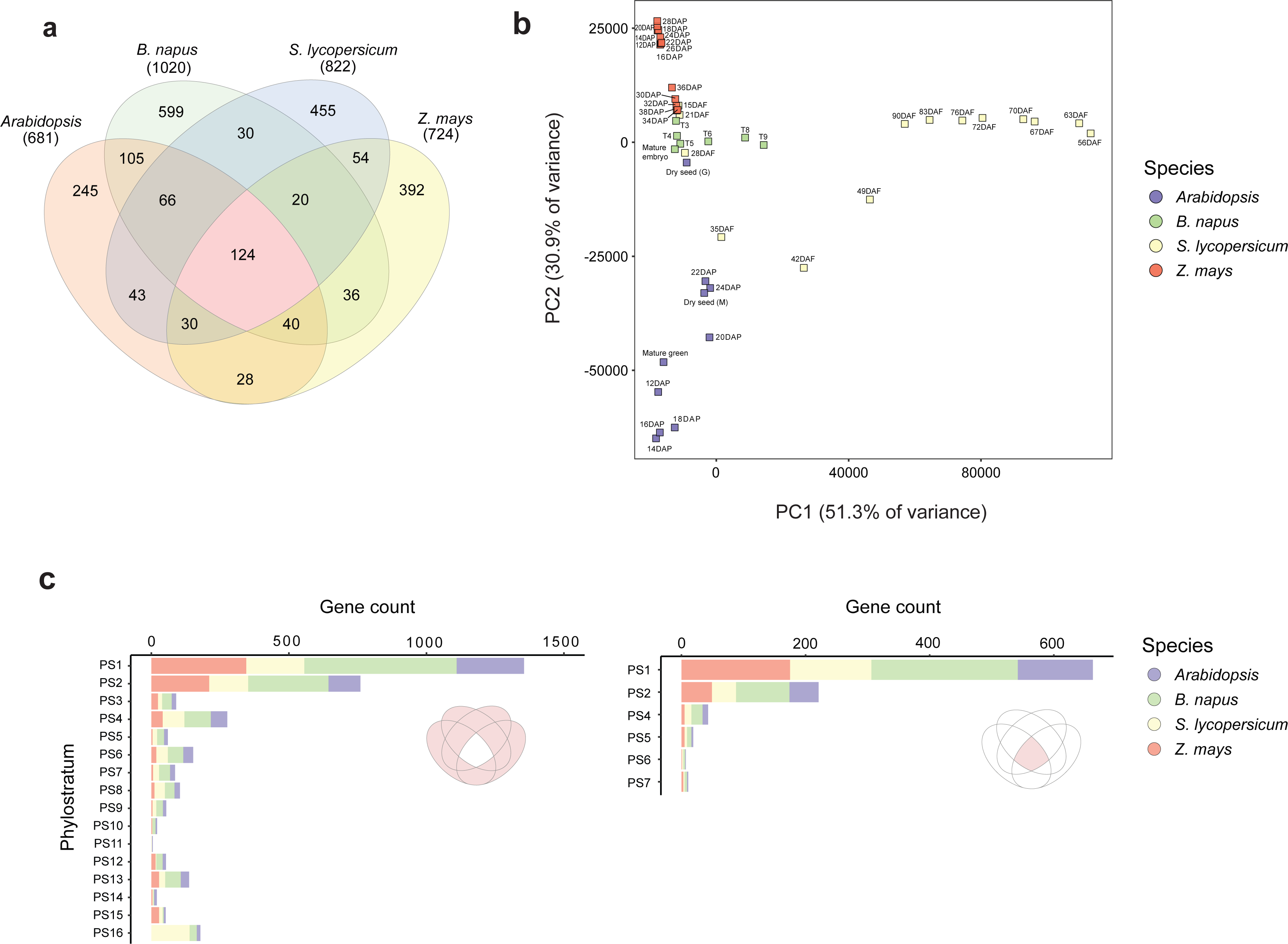
Orthogroup comparison of the top 5% maturation genes in four angiosperm species. **a**, Venn diagram showing the overlap in orthogroups that are represented within the top 5% maturation genes from *Arabidopsis*, *B. napus*, tomato (*S. lycopersicum)*, and maize (*Z. mays)*. **b**, PCA showing clustering of maturation time points from all four species based on the top 5% maturation orthogroups shown in the Venn diagram. **c**, Gene counts of orthogroups sorted according to phylostrata that are either conserved (right) or non-conserved (left) in all four species.

In summary, these findings show that the developmental reverse hourglass pattern during the seed life cycle is conserved across both dicot and monocot species. Additionally, the characteristic high TAI observed during seed maturation in these species is primarily driven by high expression of genes originating from a few relatively younger phylostrata and that are involved in both essential cellular processes and seed-specific functions.

### Embryo and endosperm tissues contribute differently to the seed maturation TAI

To obtain a better understanding of the causes underlying the high TAI profile during seed maturation we investigated the tissue-specific contribution to the overall TAI profile. The availability of tissue-specific (endosperm and embryo) RNA-seq data for a subset of the seed life cycle time points from tomato and maize facilitated this investigation (Supplemental Table 1). In tomato, we found that both endosperm and embryo contributed to the overall TAI pattern, i.e., higher TAI during mid-maturation time points were seen for both tissues (Supplemental Fig. 18). Remarkably, a noticeable difference could be seen in the pattern of a subset of the phylostrata. For instance, PS6 and PS13 had higher TAI values in the endosperm than in the embryo. On the contrary, PS9 had a relatively higher TAI value in the embryo than the endosperm (Supplemental Fig. 18). Additionally, PS16 was one of the highest contributing phylostratum in both tissues, with a more pronounced peak in the endosperm at 42 DAF.

In maize, the contrast in tissue-specific TAI contribution between endosperm and embryo was clearly discernible (Supplemental Fig. 19). The high TAI profile of PS13 observed in the whole maize seed (Fig. 2f) appeared to be almost exclusively due to the endosperm. A likely reason is that zein genes that belong to PS13, are highly expressed in the endosperm (Woo et al., 2001). To validate this finding, we recalculated the endosperm TAI for all phylostrata after removing all maize zein-annotated genes (49 genes) from the expression data (Supplemental Table 8 and Supplemental Fig. 20). Expectedly, this abolished the high TAI pattern of PS13 in the endosperm, which resulted in a pattern that was more similar to that of the embryo (Supplemental Fig. 19 and 20b).

To discern whether the stark contrast in tissue-specific contribution observed in maize was species-specific or a conserved pattern in another monocot species, we also analyzed the whole seed, embryo and endosperm RNA-seq data of barley (*Hordeum vulgare* L.) seed maturation (Kovacik et al., 2024). Phylostratigraphy of barley showed a total of 16 phylostrata (PS1-PS16, Supplemental Fig. 21a). Although the RNA-seq data only partly covered embryogenesis (4 DAP and 8 DAP) and maturation (16 DAP, 22 DAP, and 32 DAP) (Bartels et al., 1988), it still allowed us to look into the change in TAI pattern during part of barley seed life cycle. In line with the other species analyzed here, the TAI pattern of barley whole seed also increased during the transition from embryogenesis to maturation (Supplemental Fig. 21b). Moreover, we observed that three relatively young phylostrata (PS8 (Mesangiospermae), PS12 (Poaceae), and PS14 (Pooideae)) contributed more prominently to the overall TAI profile (Supplemental Fig. 22a), which aligns with what we observed in maize. Looking at the tissue-specific TAI contribution, we found that most of the contributions of PS8, PS12, and PS14 to the whole barley seed TAI pattern were derived from the endosperm (Supplemental Fig. 22b), resembling what we observed for PS13 contribution to the TAI profile in maize. The top 20 genes with the highest expression during these time points (4 DAP to 32 DAP) include, among others, genes encoding for LEA protein, defensins, glutenins, globulin, beta purothionins, alpha-amylases and gamma gliadins, and lipid transfer proteins, which are genes involved in desiccation tolerance and grain filling (Gorjanović, 2009; Yang et al., 2023) (Supplemental Table 10). Together, these findings reveal that the pronounced contribution of endosperm-expressed genes to the overall seed TAI pattern is a unifying feature of monocots, with younger PS-derived seed storage protein-related genes predominantly expressed in the endosperm.

### Functionally specialized genes drive the high TAI patterns of the seed life cycle and pollen development

“Out of the testis” is a widely accepted hypothesis concerning de novo gene evolution that pinpoints that male reproductive tissues serve as the basis for the expression of new genes (Vinckenbosch et al., 2006; Levine et al., 2006; Kaessmann, 2010). A multitude of evidence has accumulated over the years in support of this hypothesis for both animal and plant species (termed “out of the pollen” hypothesis in plants) (Betran, 2002; Begun et al., 2007; Wu et al., 2014; Cui et al., 2015; Assis, 2019; Julca et al., 2021). Given that also for pollen development, a high TAI profile has been reported (Cui et al., 2015), we investigated whether pollen development and seed maturation show expression of similar young phylostrata genes. We reanalyzed the RNA-seq data from *Arabidopsis* (ecotype Col-0) pollen development from Cui et al. (2015) to calculate the TAI values for each phylostrata. For comparisons, we also took along the egg cell data from the same transcriptome as a reference for the female gamete. The TAI values showed a progressive increase during stages of pollen development and reached a maximum value in the pollen tube cells, while the egg cell TAI value was strikingly lower (Fig. 4a). The overall pattern deviated significantly from a flat line (*p_flt_* = 4.38X10^-14^), indicating the presence of an evolutionary signature. We also found that several younger phylostrata highly contributed to the TAI during pollen development (Fig. 4b and Supplemental Fig. 23), which was expected based on previous reports (Cui et al., 2015; Gossmann et al., 2016b; Julca et al., 2021).

**Fig. 4.**
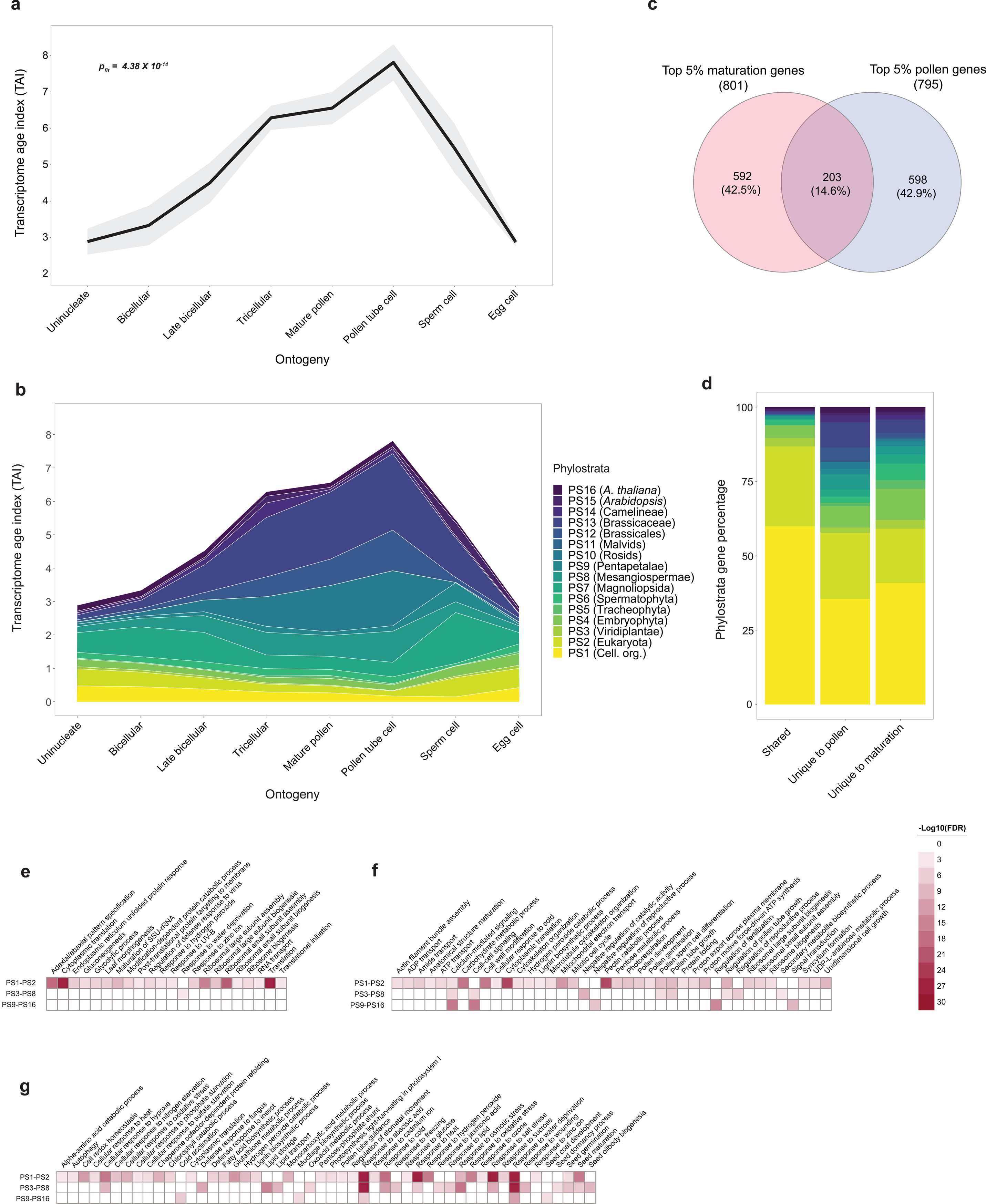
Comparison of the phylotranscriptome of Arabidopsis pollen development and seed maturation. **a**, TAI pattern during *Arabidopsis* pollen development and egg cell stage. The p-value indicates the significance of deviation from a flatline test. Shaded regions around the TAI line indicates the standard deviation (s.d.) calculated from 1000 permutations. **b**, Contribution of individual phylostratum to the overall TAI profile. **c**, Venn diagram showing the overlap between top 5% genes during seed maturation and pollen development. **d**, Relative percentage of genes from each phylostrata comprising genes that are shared and unique between the seed maturation and pollen development. **e-f**, Heatmap showing GO terms that are enriched in genes that are shared, unique to pollen development, and unique to seed maturation, respectively. The color indicates the -log10(FDR) value between a range from 0 to 30.

We then questioned whether similar genes underlay the high TAI profile during pollen development and seed maturation in *Arabidopsis*. To examine this further, we extracted the top 5% genes with the highest average expression during pollen development (uninucleate until sperm cells, 795 genes) (Supplemental Table 11). Similar to what we observed for seed life cycle TAI (Supplemental Fig. 4c), removing the top 5% expressed genes from the pollen dataset largely decreased the high TAI profile (Supplemental Fig. 24). Comparison of the top 5% pollen genes with the top 5% seed maturation expressed genes (Supplemental Table 6) showed an overlap of 203 genes, with 592 and 598 genes unique to pollen development and seed maturation, respectively (Fig. 4c). We also found that over 80% of the shared genes belonged to the older phylostrata (PS1 and PS2) (Fig. 4d). In contrast, a higher percentage of younger phylostrata genes was observed among the unique genes for both pollen development and seed maturation (Fig. 4d). A GO enrichment further confirmed that shared genes between both tissues were mostly from associated with primary cellular processes such as translation, mRNA maturation, and energy metabolism (Fig. 4e). On the contrary, highly expressed genes specific either to pollen development or seed maturation were enriched GO terms, which are related to the functions specific to each respective organ (Fig. 4e-g, respectively). For example, enrichment of negative regulation of the reproductive process, regulation of fertilization, and signal transduction were uniquely observed for younger phylostrata genes (PS9-PS16) for pollen, while the same PS were specifically enriched in genes involved in cold acclimation, mucilage biosynthetic process, and seed coat development for seed maturation. These findings show that, while pollen and seed life cycle share a similar reverse hourglass, the genes driving these patterns are functionally specialized for each structure.

## Discussion

One of the major applications of phylotranscriptomics in animals and plants has been in understanding the gene evolution underlying the molecular landscape of embryo development across species. With the application of phylotranscriptomic indices (TAI and TDI), biologists have been able to provide molecular support for the hourglass-like pattern of animal embryo development, highlighting deep morphological and genetic conservation at mid-embryonic stages, i.e., the phylotypic stage (Richardson, 1995; Liu et al., 2021). However, a reverse hourglass-like development was observed during embryogenesis in three animals with spiralian development (Wu et al., 2019), highlighting mid-embryonic development as a divergent phase in spiralians. This finding challenged the conventional hourglass model of embryo development and suggests that different evolutionary mechanisms may shape development in different lineages.

In plants, an hourglass-like development has been reported for embryonic (embryogenesis) and also for post-embryonic (germination) development using the model plant *Arabidopsis* (Quint et al., 2012; Drost et al., 2016). Here, we analyzed the phylotranscriptome throughout the seed life cycle in angiosperms to understand evolutionary patterns across key phase transitions during seed development. This approach bridges the gap between embryogenesis and germination by incorporating recent high-resolution transcriptomic data from seed maturation (Artur et al., 2024). We show that *Arabidopsis* seed maturation exhibits a relatively younger transcriptome with higher TAI and TDI values in relation to embryogenesis and germination. From a broader perspective, the Arabidopsis seed life cycle resembles a reverse hourglass pattern, with the transcriptome of early and late developmental phases (embryogenesis and germination, respectively) highly expressing older genes, while the mid-developmental phase (maturation) exhibits a more diverse transcriptome with a higher proportion of younger genes (Fig. 5). We also observed that genes originating from intermediate and younger phylostrata represented a higher percentage of the top-expressed genes during maturation compared to the adjacent phases. Such genes are specifically enriched for biological functions related to seed maturation-related traits such as seed storage reserve, seed coat development, germination, dormancy, and stress response.

**Fig. 5.**
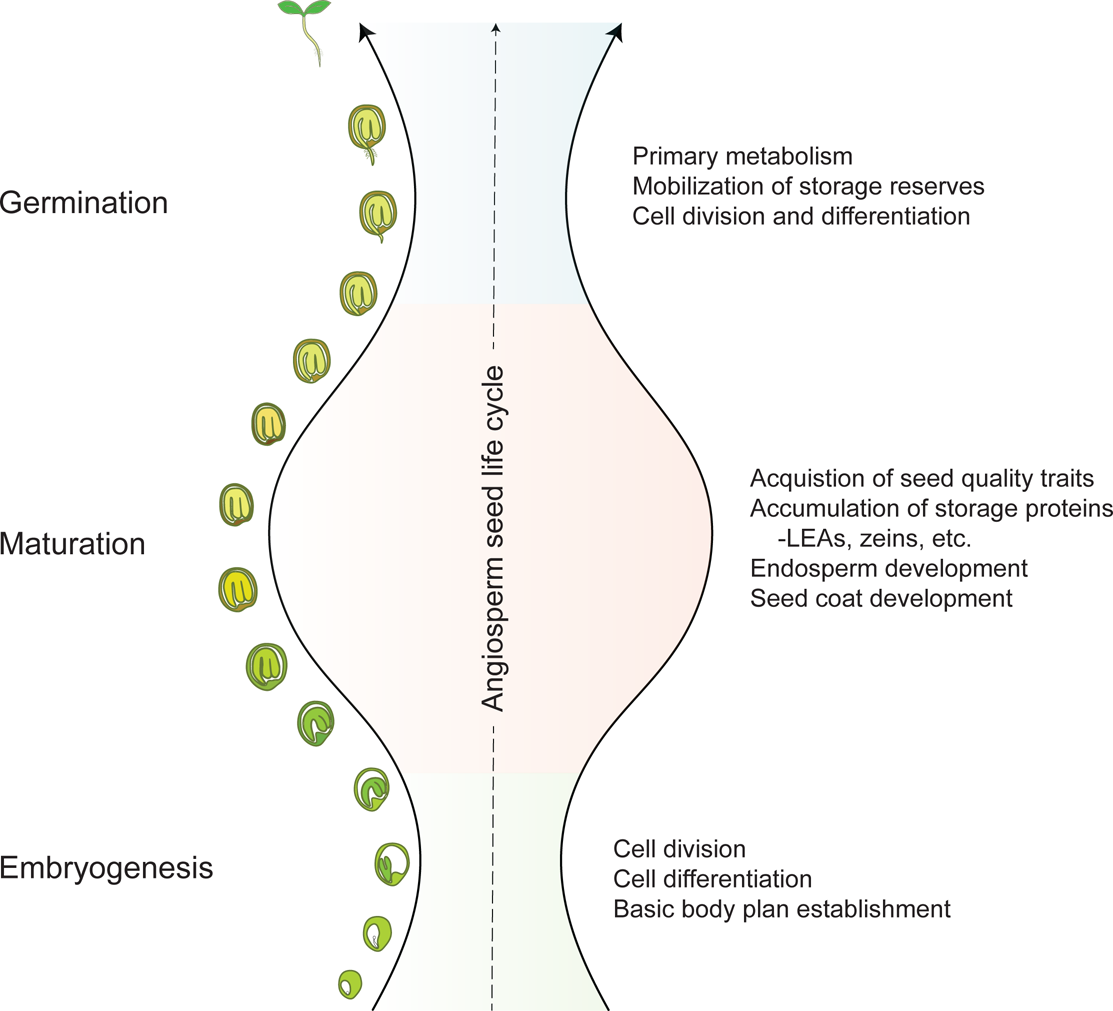
Developmental reverse hourglass model of angiosperm seed life cycle. Schematic presentation of the reverse hourglass model of the seed life cycle indicating the mid-development phase (maturation) characterized by expression of younger genes involved in seed-specific traits, while during the flanking phases (embryogenesis and germination) expression of older genes involved in basic cellular processes is more dominant.

Reverse hourglass-like relationships have also been previously reported while comparing the embryonic development between species in both animals and plants. A recent example of this is the comparison between maize and *Physcomitrium patens* embryo development (Wu et al., 2024). While each species showed an hourglass embryonic development, interspecific comparisons showed that the transcriptome of the phylotypic stage is more divergent between the two species than in earlier and later developmental stages. A similar observation was also made for inflorescence development in maize and sorghum (Leiboff and Hake, 2019). Here, we compared the TAI pattern of phases of the seed life cycle in *Arabidopsis*, *B. napus*, tomato, and maize and we found broader conservation of the reverse hourglass development. Despite conservation in the overall pattern, the genes underlying the mid-developmental phase, i.e. maturation, were divergent among different species. Several morpho-physiological seed traits, such as the type of storage proteins, seed coat composition, endosperm differentiation, and acquisition stress-resilience characteristics are obtained during maturation, and seeds from different species and ecotypes exhibit high diversity in these traits (Sreenivasulu and Wobus, 2013; Leprince et al., 2016; Saatkamp et al., 2019). Therefore, a conserved reverse hourglass pattern is a feature that may have facilitated the evolution and diversification of seed traits amongst angiosperm lineages.

In most angiosperm seeds (termed ‘orthodox’ seeds), a desiccation phase occurs during later stages of maturation and, therefore, these seeds need to acquire desiccation tolerance before reaching a critical moisture level (Leprince et al., 2016; Smolikova et al., 2020). Thus, osmotic stress is an inherent feature of orthodox seed development. Our findings showed that biological processes related to ABA response and water deprivation were enriched in the top 5% of maturation-expressed genes across all three phylostrata age groups (PS1-PS2, PS3-PS8, and PS9-PS16). This suggests the contribution of both old and young genes towards the evolution of osmotic stress tolerance during seed maturation. Amongst the most well-known examples of desiccation-associated genes, *LEAs* are group with distinct evolutionary histories that are highly expressed during seed maturation (Hundertmark and Hincha, 2008; Artur et al., 2019). In our phylotranscriptome analysis we found several *LEA*s amongst the most highly expressed genes during both *Arabidopsis* and maize seed maturation. Remarkably, the maturation-expressed LEAs from *Arabidopsis* derived from a broad range of phylostrata (spanning from PS1 to PS14). Despite their undeniable association with desiccation stress, several questions still remain about the relationship between LEAs evolution and their structural diversity and molecular functions in response to cellular water loss (Hernández-Sánchez et al., 2022). Our finding suggests that the combined high expression of *LEA* genes from diverse origins underlies seed maturation and might be a requirement for seed desiccation tolerance acquisition in seeds.

The association of genes of younger origin with overall stress responses have been described for multiple species. For example, lineage-specific genes of *de novo* origin in *Arabidopsis* have been shown to be enriched for stress responsiveness (Donoghue et al., 2011). Similarly, lineage-specific orphan genes in sugarcane (*Saccharum* spp.) are enriched for external stimulus and defense response and are more responsive to cold and osmotic stress (Cardoso-Silva et al., 2022). A comparison of genomes from multiple rice species showed that Poaceae and Oryzeae-specific genes with functional domains are enriched for stress and defense-related functions (Stein et al., 2018). Similar associations have also been reported for young genes from more distant branches in the tree of life. For instance, in yeasts, *de novo* genes arising from previously non-coding genomic regions were ascribed to stress-related functionality (Carvunis et al., 2012). Moreover, a recent study highlighted a link between stress response and gene age in yeast, where the cells were more likely to express younger genes under stress conditions, leading to the functionalization of recently emerged genes (Doughty et al., 2020). In addition, a number of *de novo* genes in yeast were only found to be expressed under oxidative stress in contrast to control conditions (Blevins et al., 2021). In *Daphnia*, lineage-specific genes were found to be enriched in transcriptome datasets generated under ecologically challenging conditions (Colbourne et al., 2011). Thus, our finding that biological processes related to desiccation and osmotic stress signaling were enriched in young phylostrata genes during *Arabidopsis* maturation further supports the well-documented association of young genes with overall stress responses.

Our tissue-specific phylotranscriptome analysis revealed that embryo and endosperm tissues contribute differently to the phylotranscriptome pattern of seed maturation. Notable was the near-exclusive contribution of the endosperm towards a high seed maturation TAI consistently found in two monocots (maize and barley). This could be explained by the persistent nature of the endosperm in monocots (Olsen, 2004; Sreenivasulu et al., 2010; Sreenivasulu and Wobus, 2013). On the contrary, in dicots, after cellularization, the embryo gradually absorbs the endosperm during maturation (Berger, 1999; Berger et al., 2006; Sreenivasulu and Wobus, 2013). As a result, the endosperm is the major compartment for protein storage and nutrient reserves in monocots, while in dicots, they are mainly localized in the cotyledons (Sreenivasulu and Wobus, 2013). One of the key factors underlying the reverse hourglass pattern in the monocots addressed here is the diverse range of storage protein-associated genes coming from several younger phylostrata. Therefore, it is highly likely that the different sources of high TAI during seed maturation explains the differences in endosperm developmental, physiological, and molecular characteristics in both monocots and dicots.

*De novo* genes are well known to be expressed in male reproductive tissues (Betran, 2002; Levine et al., 2006; Kaessmann, 2010). Even within angiosperms, multiple studies have shown that younger genes (also those of *de novo* origin) are more prone to be expressed in the male gamete, i.e., pollens (Williams, 2008; Wu et al., 2014; Cui et al., 2015; Gossmann et al., 2016a). By comparing the phylotranscriptomic patterns of both *Arabidopsis* pollen development and seed maturation we observed a higher TAI owing to high expression of multiple young phylostrata genes. A cross-comparison of the top expressed genes in both structures showed a partial overlap, mostly consisting of genes from older phylostrata involved in basic cellular functions. Despite the similar reliance on young genes, both stages of development seem to express a unique set of genes that are functionally specialized to the respective developmental stage. Due to the relevance of pollens and seeds in natural selection and in the evolution of angiosperms, these new genes may get fixed by gaining specialized functions related to either of the two developmental stages (Stanton et al., 1986; Moles et al., 2005; Williams, 2008; Sims, 2012; Coen and Magnani, 2018).

Together, our findings suggest that, similar to the ‘out of the pollen’ hypothesis for the evolution of novel genes during pollen development (Cui et al., 2015; Gossmann et al., 2016a), seed maturation might also serve as a landscape that offers similar opportunities to test new genes and facilitate functional specialization.

## Materials and Methods

### Transcriptome datasets and RNA-seq analysis

Transcriptome datasets covering the three phases of the seed life cycle (embryogenesis, maturation, and germination) were obtained from publicly available RNA-seq libraries from previous studies (Supplemental Table 1). For *Arabidopsis* and *B. napus*, samples until the bent cotyledon stage were considered part of embryogenesis (Baud et al., 2002; Bianchetti et al., 2024) and subsequent time points until the dry seed stage were considered part of maturation. For *B. napus*, the RNA-seq time points are expressed as thermal time (T1-T6 and T8-T9) where each time point represents growing degree days (GDD) from the start of flowering at a base temperature of 0^0^ C (detailed in Bianchetti et al., 2021). T1-T2 correspond to the transition from torpedo to bent cotyledon stages, T2-T6 to seed filling, and T6-T9 represents the final stages of seed maturation i.e., late maturation (Bianchetti et al., 2021). Based on the onset of seed filling and maturation traits, the tomato RNA-seq dataset was divided into two parts: 15 days after flowering (DAF) to 21 DAF indicating the last stages of embryogenesis, and 28 DAF onwards corresponding to maturation (Bizouerne et al., 2021). The maize transcriptome was grouped into embryogenesis (0 days after pollination (DAP) to 10 DAP) and maturation (12 DAP to 38 DAP) based on the expression of zein genes which were considered as indicators of seed filling (Chen et al., 2014). As for barley, time points 4 DAP and 8 DAP were considered part of embryogenesis while 16 DAP to 32 DAP were considered as part of maturation based on initiation of seed filling (Bartels et al., 1988). Fastq files were downloaded from the National Center for Biotechnology Information (NCBI) using the Sequence Read Archive (SRA) Toolkit (v3.0.3) and mapped to their respective genomes (indicated in Supplemental Table 1) using the nf-core (Di Tommaso et al., 2017; Ewels et al., 2020) rnaseq pipeline (v3.9). The pipeline uses bedtools (v2.30.0) (Quinlan and Hall, 2010), bioconductor-summarizedexperiment (v120.0) (Martin Morgan, 2017), bioconductor-tximeta (v1.8.0) (Love et al., 2020), fastqc (v0.11.9), gffread (v0.12.1) (Pertea and Pertea, 2020), picard (v2.27.4), star (v2.7.10a) (Dobin et al., 2013), salmon (v1.5.2) (Patro et al., 2017), stringtie (v2.2.1) (Pertea et al., 2015), Trimgalore (v0.6.7), and ucsc (v377). The resulting gene transcript per million (TPM) values were used for all downstream analysis.

### Phylostratigraphy and TAI calculation

The phylostratigraphy of all plant species was constructed using GenEra (v1.4.0) (Barrera-Redondo et al., 2023) with default parameters. All genes of a given target genome were queried against a custom reference protein database comprising of 4,940 proteomes using DIAMOND (Buchfink et al., 2015) blastp in more-sensitive mode with an e-value threshold of 10^-5^. The custom database comprised of proteomes of 175 plants, 582 fungi, 391 vertebrates, 384 invertebrates, 96 protozoa, 1,055 archaea, and 2,257 bacteria (Supplemental Table 2). Blastp hits were considered as homologs of the query protein and were used to determine gene family founder events and relative age of a gene.

TAI of RNA-seq samples were calculated as described previously (Domazet-Lošo and Tautz, 2010) using the myTAI (v0.9.3) (Drost et al., 2015) package in R (v4.3.2). TAI values were calculated as the weighted mean of phylostratum rank (*ps_i_*) of a given gene *i* by the expression level (*e_is_*) in the transcriptome of an RNA-seq sample *s*,

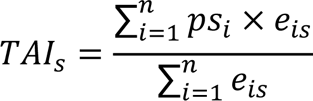

Where *n* represents the total number of genes present in the RNA-seq sample *s*. A high TAI value indicates a younger transcriptome age, whereas a lower TAI corresponds to a more ancient transcriptome.

### Ka/Ks and TDI calculation

The ratio of synonymous substitution rate and non-synonymous substitution rate (Ka/Ks) for *A. thaliana* and *A. lyrata* orthologous gene pairs were determined using the R package orthologr (Drost et al., 2015). The following arguments were used for Ka/Ks calculation, ortho_detection = “RBH”, aa_aln_type = “pairwise”, aa_aln_tool = “NW”, codon_aln_tool = “pal2nal”, dnds_est.method = “Comeron”. Gene pairs with a Ka/Ks < 2 were retained. Based on the Ka/Ks values, *Arabidopsis* genes were binned into ten divergence strata (DS) from DS1-DS10 (low to high). The TDI was calculated for the same RNA-seq samples by replacing the PS with the DS, as shown below,

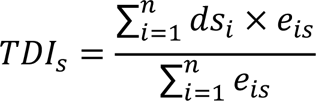

A high/low TDI value indicates a conserved/divergent transcriptome. Similarly, the Ka/Ks ratio was calculated for *B. napus*, tomato, and maize genes based on ortholog comparison with *B. oleracea*, *S. pimpinellifolium*, and *Z. diploperennis* genomes, respectively. Subsequently, the TDI values were calculated using the Ka/Ks ratio.

### Reverse hourglass test and individual phylostrata contribution

The reverse hourglass test was carried out using the myTAI package. Each species’ developmental stages were partitioned into three groups – early, mid, and late. For *Arabidopsis* and *B. napus*, embryogenesis time points were grouped as early, maturation time points as mid, and germination time points as late (Supplemental Table 1). On the other hand, the tomato RNA-seq dataset comprised of time points from late embryogenesis (15 DAF and 21 DAF) until the end of seed maturation (90 DAF). In this case, we considered 15 DAF to 21 DAF as early, 35 DAF to 70 DAF as mid and 72 DAF to 90 DAF as late stages. In case of tomato and maize, late maturation time points were grouped as late stages. Similarly, for maize, 0 to 8 DAP was considered as early, 10 DAP to 28 DAP as mid, and 30 DAP to 38 DAP as late stages.

Relative expression and individual phylostrata contribution were also calculated with the myTAI package in R. The data was plotted using ggplot2 (v3.5.0) (Wickham, 2009).

### Determining top maturation expressed genes

To determine the top maturation-expressed genes with the highest expression, we calculated the mean TPM expression for each gene over maturation time points (Supplemental Table 1). Genes with an average TPM of ≥ 1 were taken along and sorted based on average expression from high to low. From this list, the top 5% of genes were considered the top maturation-expressed genes.

### Gene ontology (GO) enrichment analysis

*A. thaliana* GO slim terms were downloaded from TAIR (https://www.arabidopsis.org/). GO terms for *B. napus*, tomato, and maize proteins were derived from eggNOG-mapper (v2) (Huerta-Cepas et al., 2019; Cantalapiedra et al., 2021) using default parameters. GO enrichment was carried out using the R package topGO (v2.54.0) (Adrian Alexa, 2023) using Fisher’s exact test. GO terms with a false discovery rate (FDR) of ≤ 0.001 were retained. The list was sorted based on -log10(FDR) from highest to lowest, and only the top 15 non-redundant GO terms were shown from each PS group.

### Orthogroup and interspecies maturation transcriptome comparison

Protein sequences of all five species in this study were used to determine orthologous gene groups (orthogroups) using Orthofinder (v2.5.5) (Emms and Kelly, 2019).

To perform interspecies maturation transcriptome comparisons, we first filtered the transcriptome of each species to retain genes that have a TPM expression of 2 in at least 3 RNA-seq samples. Next, we converted the gene identifiers (IDs) to their respective orthogroups and averaged the gene expression per orthogroup. Subsequently, we combined the orthogroup average expression from the four species – *A. thaliana*, *B. napus*, tomato, and maize to retain transcriptome information for orthogroups shared between all species. This data was used to create a principal component analysis (PCA) plot to show the distribution of the RNA-seq samples for all four species along different stages of seed life cycle.

## Data availability

All RNA-seq used in this study are detailed in Supplemental Table 1.

## Acknowledgements

We thank Dr. Kin Pan Chung for the critical feedback in the written manuscript. This work was supported by The Netherlands Organization for Scientific Research (NWO), NWO-ENW Veni (project Fine Drying VI.Veni.202.038) to M.A.S.A. and NWO VICI (project Seeds4Ever 17047) to L.B..

## Author Contributions

A.A.S. and M.A.S.A. designed the project. A.A.S. analysed the data. A.A.S. prepared the figures and tables. A.A.S. drafted the manuscript. A.A.S., M.A.S.A. and L.B. edited and revised the manuscript.

## Competing Interests

The authors declare no competing interests.

## Supplemental Information

### Supplemental Tables Legends

**Supplemental Table 1.** RNA-seq datasets of the different species used in the study along with their sources.

**Supplemental Table 2.** List of genomes used for determining phylostratiography of plant species used in the study. All genomes were downloaded from the NCBI refseq database.

**Supplemental Table 3.** Phylostratiography of Arabidopsis genes determined using GenEra (v1.4.0).

**Supplemental Table 4.** Ka/Ks ratio of *A. thaliana* genes as compared to *A. lyrata*.

**Supplemental Table 5.** Arabidopsis maturation expressed genes sorted based on corresponding phylostrata.

**Supplemental Table 6.** Top 20% Arabidopsis genes with the highest average expression during maturation time points. Genes were sorted based on average expression (TPM). From this list the top 160, 320, 801, 1602, and 3203 genes were used as top 1%, 2%, 5%, 10%, and 20% in the analysis shown in Supplemental Fig. 4. The top 5% genes are shown in bold.

**Supplemental Table 7.** Phylostratiography of *B. napus* (Bna), maize (*S. lycopersicum,* Sly), maize (*Z. mays,* Zma), and *H. vulgare* (Hvu) determined using GenEra.

**Supplemental Table 8.** List of zein encoding genes in the maize genome (B73). All zein genes belong to phylostrata 13 (Panicoideae).

**Supplemental Table 9.** List of top 5% genes with the highest average expression (TPM) during seed maturation in the three species - B. napus, S. lycopersicum, and Z. mays.

**Supplemental Table 10.** *H. vulgare* genes expressed during parts of the seed life cycle sorted from high to low based on average expression (TPM).

**Supplemental Table 11.** Top 5% genes with highest average expression (TPM) during pollen development in Arabidopsis.

**Supplemental Table 12.** List of LEA genes expressed during seed maturation in Arabidopsis.

---

**Supplemental Fig. 1.**
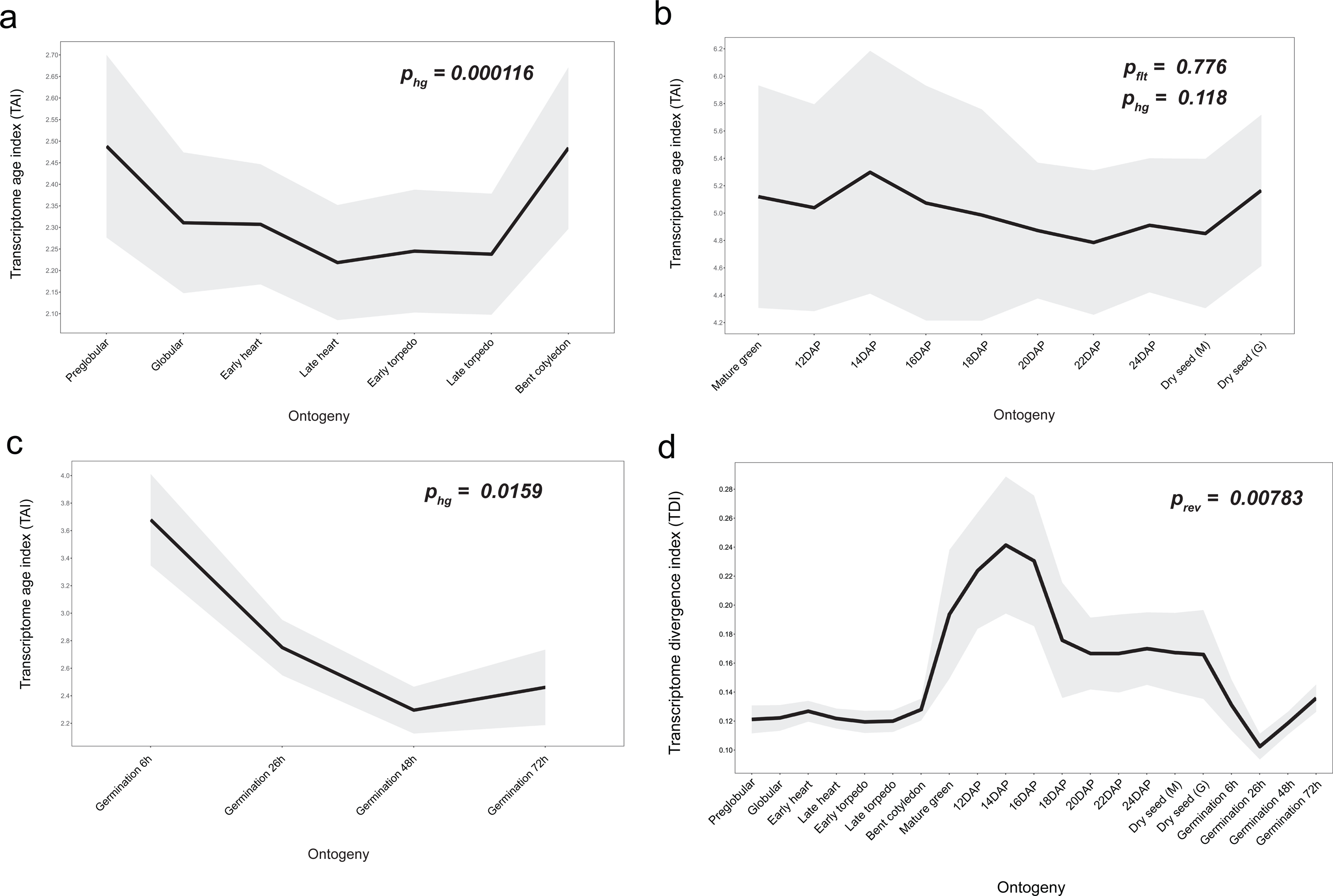
Phylotranscriptomic pattern during individual phases of the *Arabidopsis* seed life cycle - **a**, embryogenesis; **b**, maturation; and **c**, germination. P-values are shown as *p_hg_* and *p_flt_* to indicate the significance of an hourglass and flat-line test. d, TDI pattern during seed life cycle in *Arabidopsis*. The overall pattern significantly resembles a reverse hourglass pattern.

**Supplemental Fig. 2.**
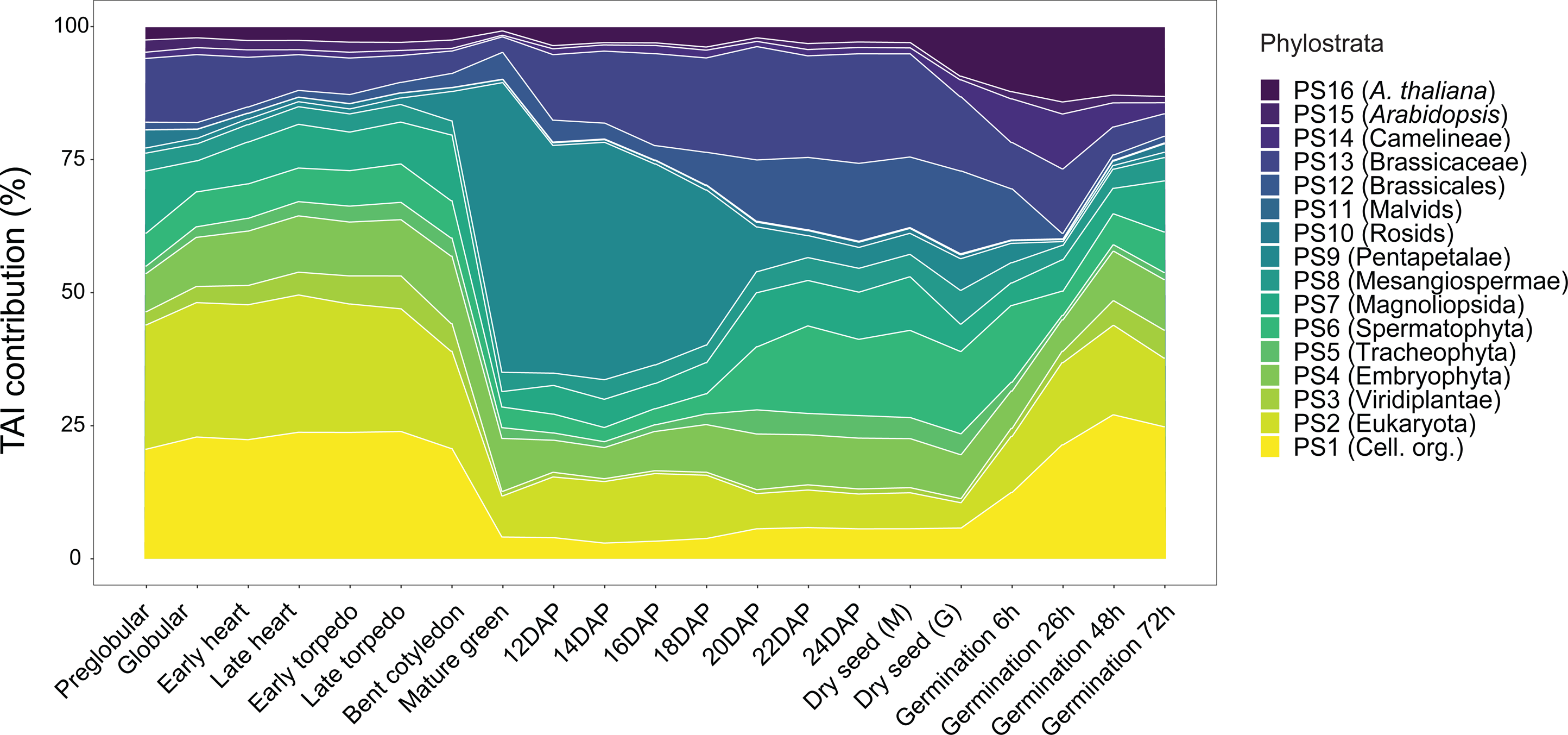
Percentage contribution of each phylostrata (PS1-PS16) to the overall TAI profile during the *Arabidopsis* seed life cycle.

**Supplemental Fig. 3.**
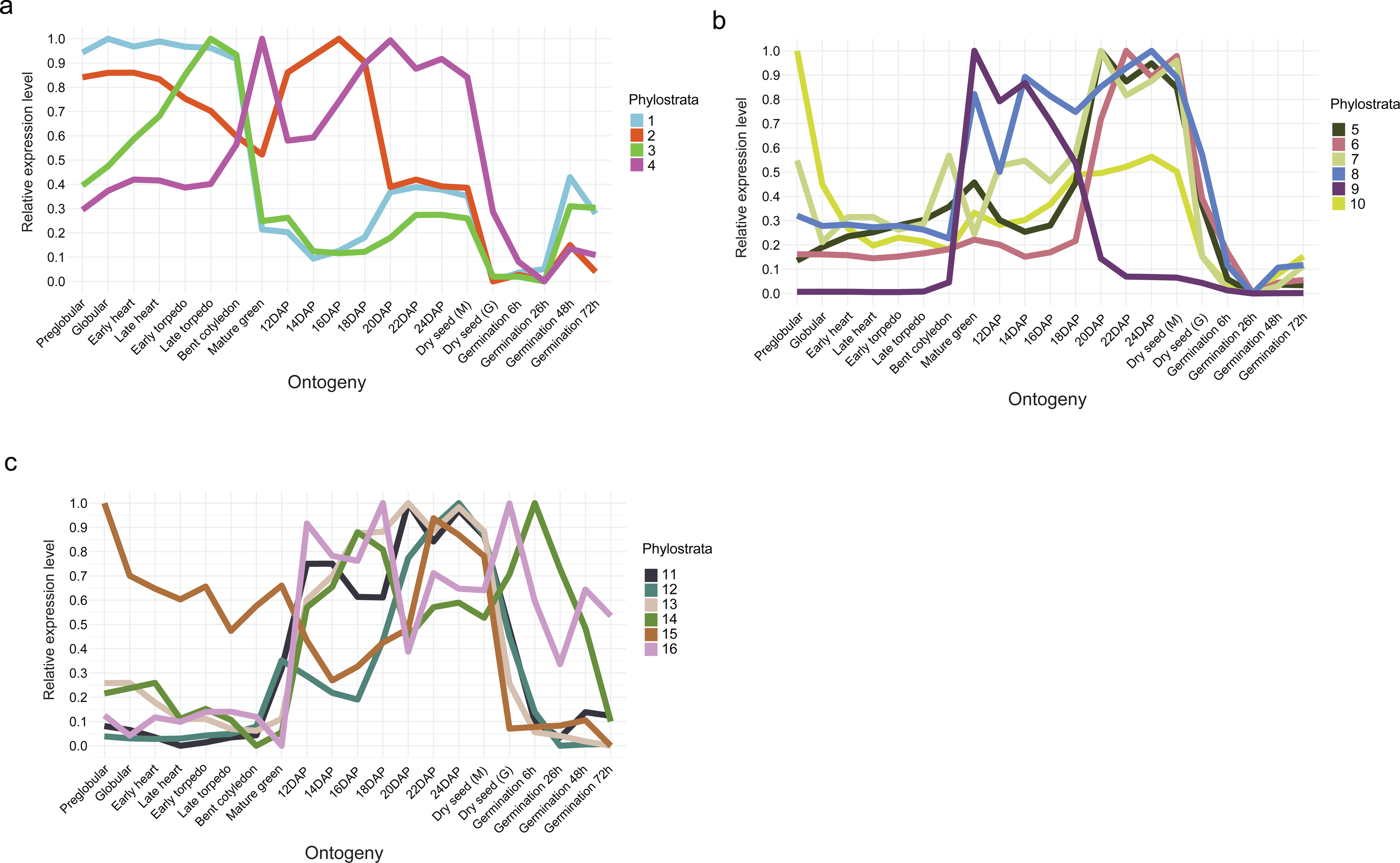
Relative expression of individual phylostrata during *Arabidopsis* seed life cycle. **a**, PS1-PS4; **b**, PS5-PS10; and **c**, PS11-PS16.

**Supplemental Fig. 4.**
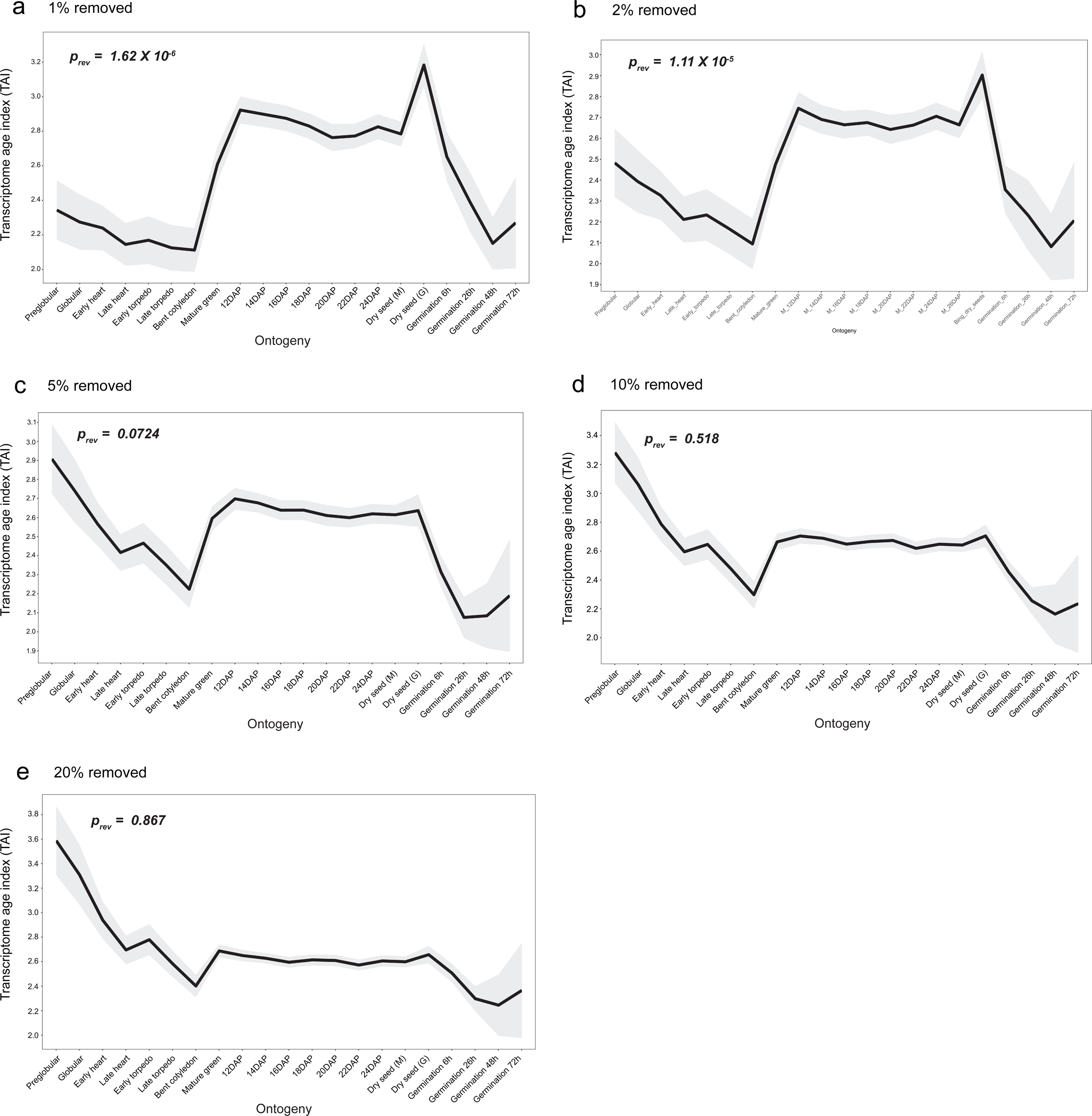
TAI profile over *Arabidopsis* seed life cycle after removing the top a, 1%; b, 2%; c, 5%; d, 10%; or e, 20% highly expressed genes during seed maturation (Mature green stage to Bing Dry seed).

**Supplemental Fig. 5.**
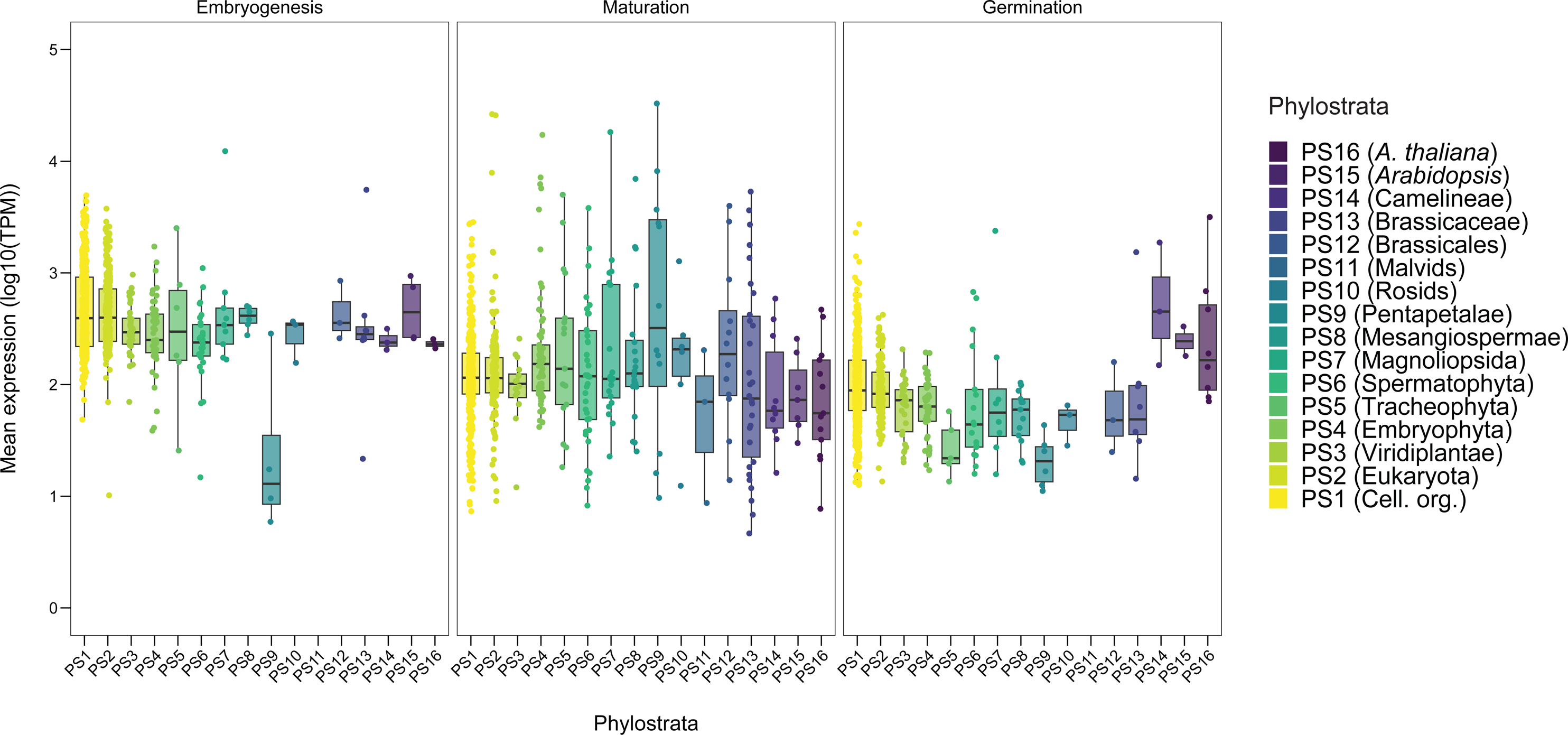
Mean expression (log10-TPM) of the top 5% genes with the highest average expression during each of the three phases of the seed life cycle. From left to right – top 5% embryogenesis expressed genes, top 5% maturation expressed genes, and top 5% germination expressed genes. The mean expression is sorted based on phylostrata.

**Supplemental Fig. 6.**
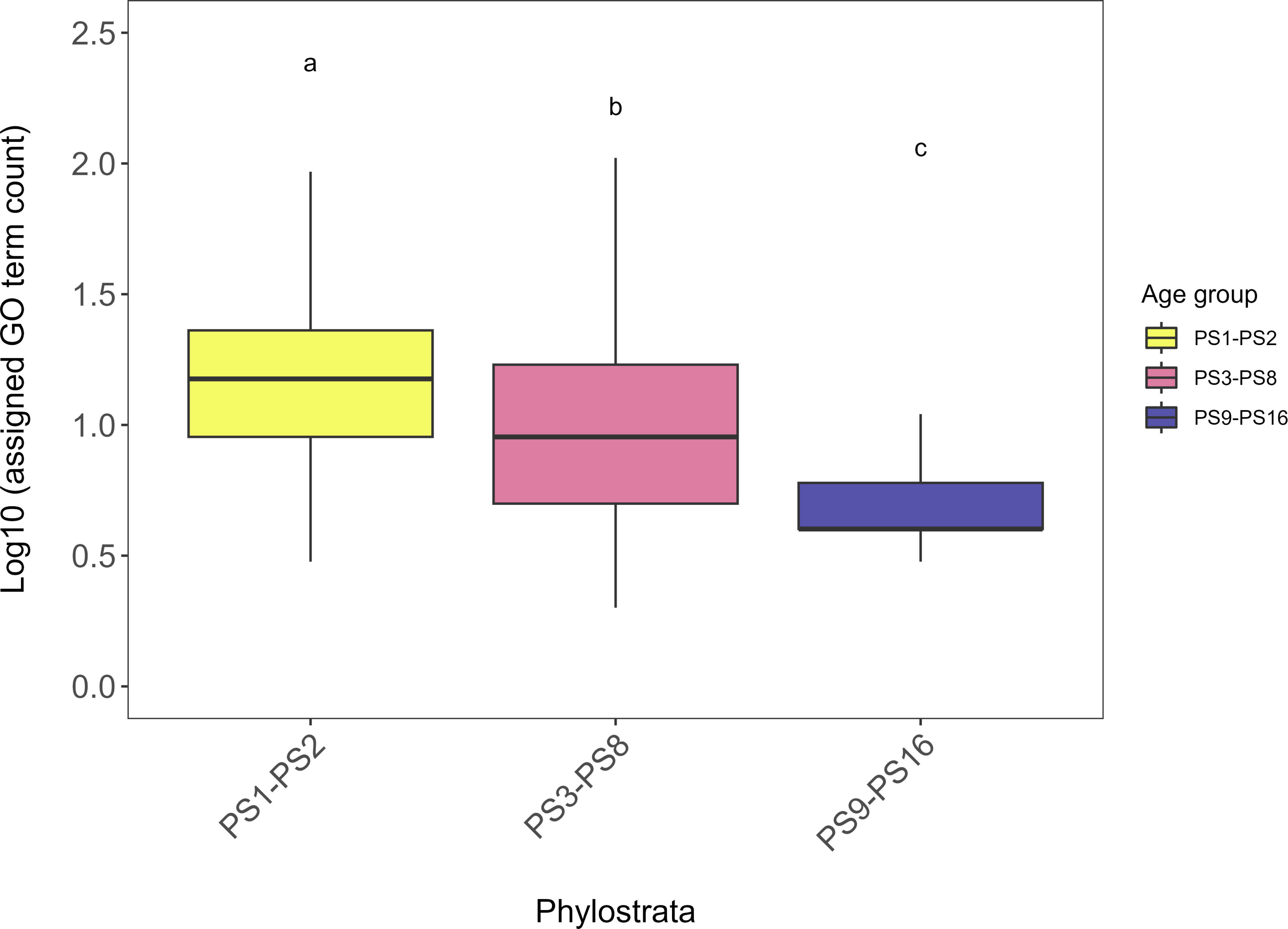
Number of GO annotation terms assigned to *Arabidopsis* genes from different phylostrata groups. Younger phylostrata groups (PS3-PS8 and PS9-PS16) have significantly (Kruskal-Wallis test, *p-value* < 0.001) fewer annotation terms than older phylostrata groups (PS1-PS2).

**Supplemental Fig. 7.**
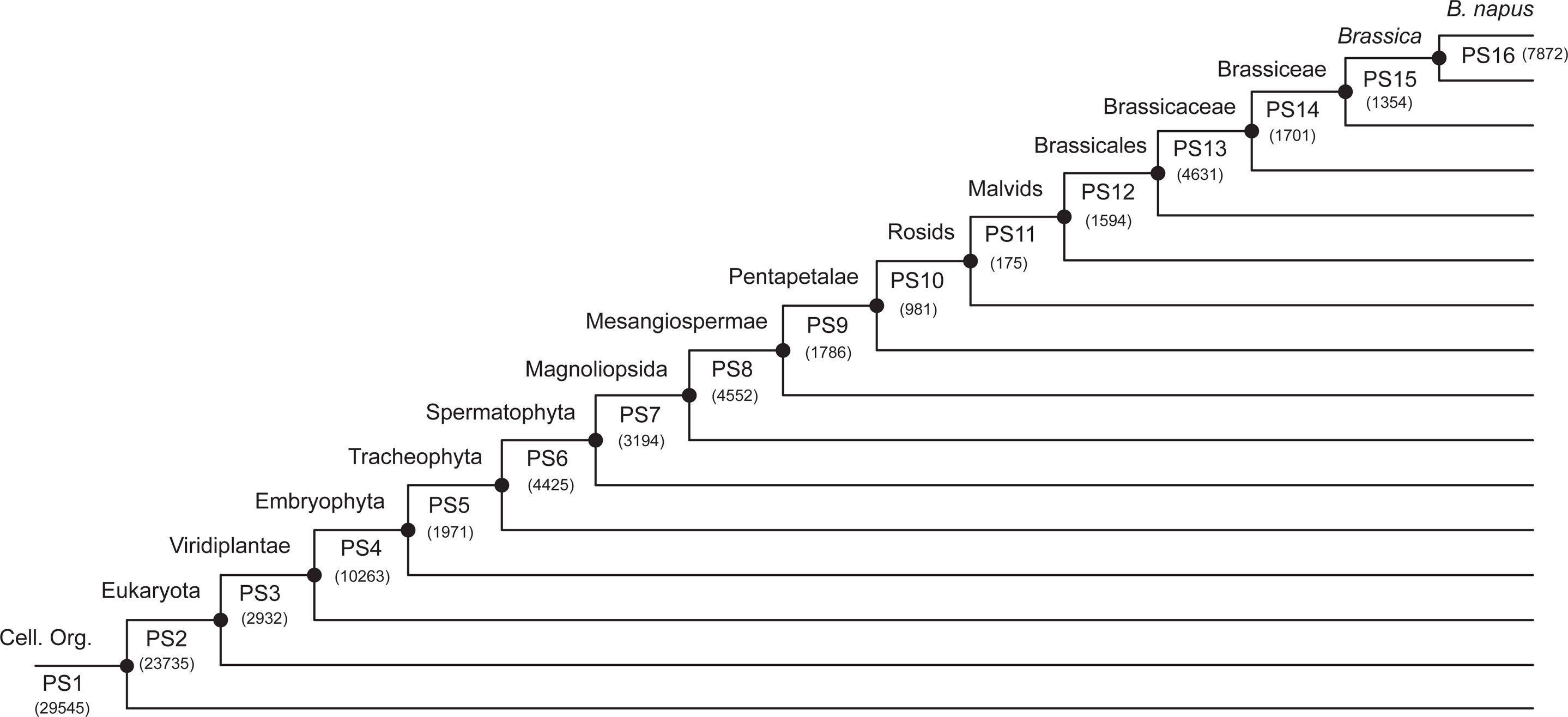
Phylostratigraphy of *B. napus* genes.

**Supplemental Fig. 8.**
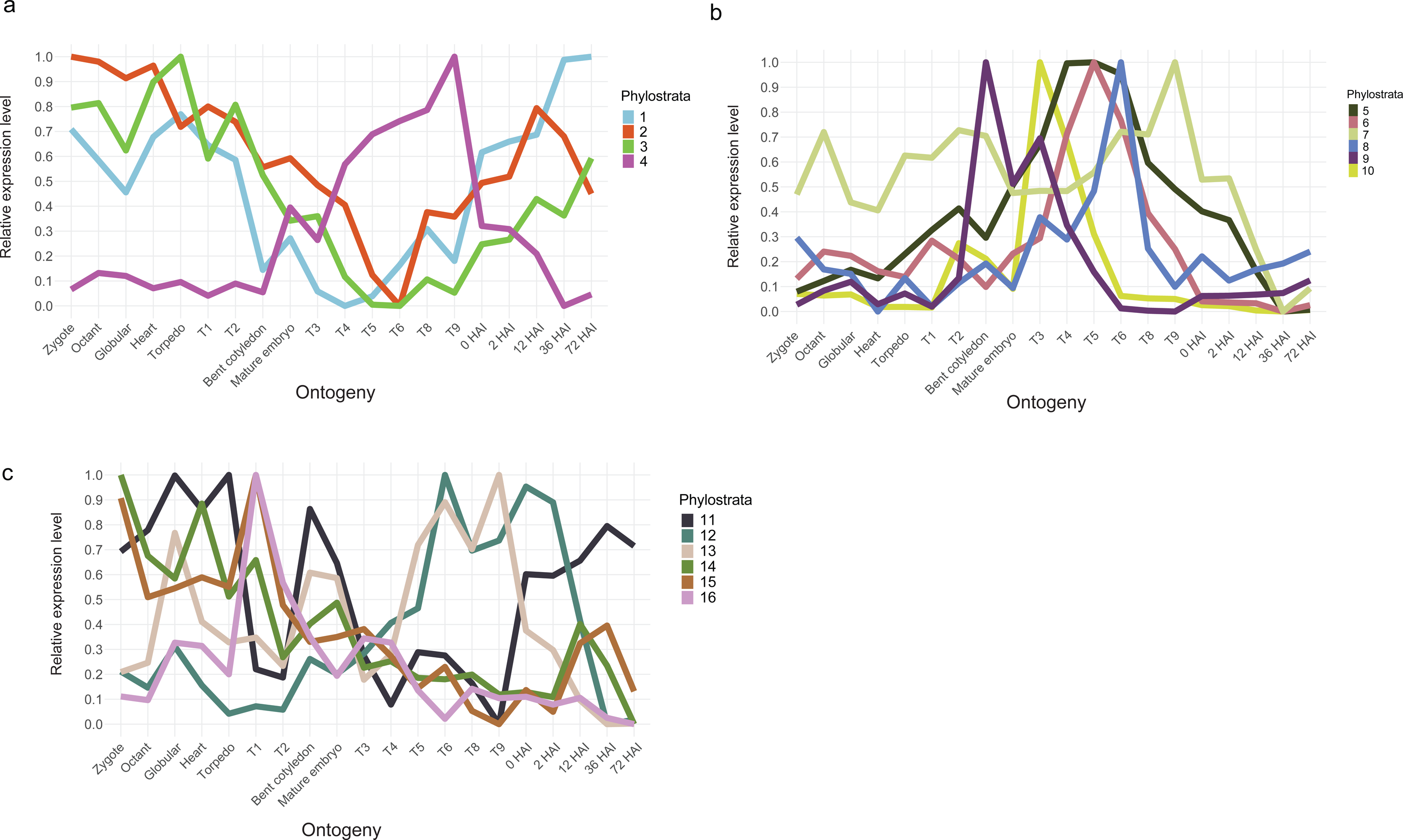
Relative expression of each phylostrata during *B. napus* seed life cycle. **a**, PS1-PS4; **b**, PS5-PS10; and **c**, PS11-PS16.

**Supplemental Fig. 9.**
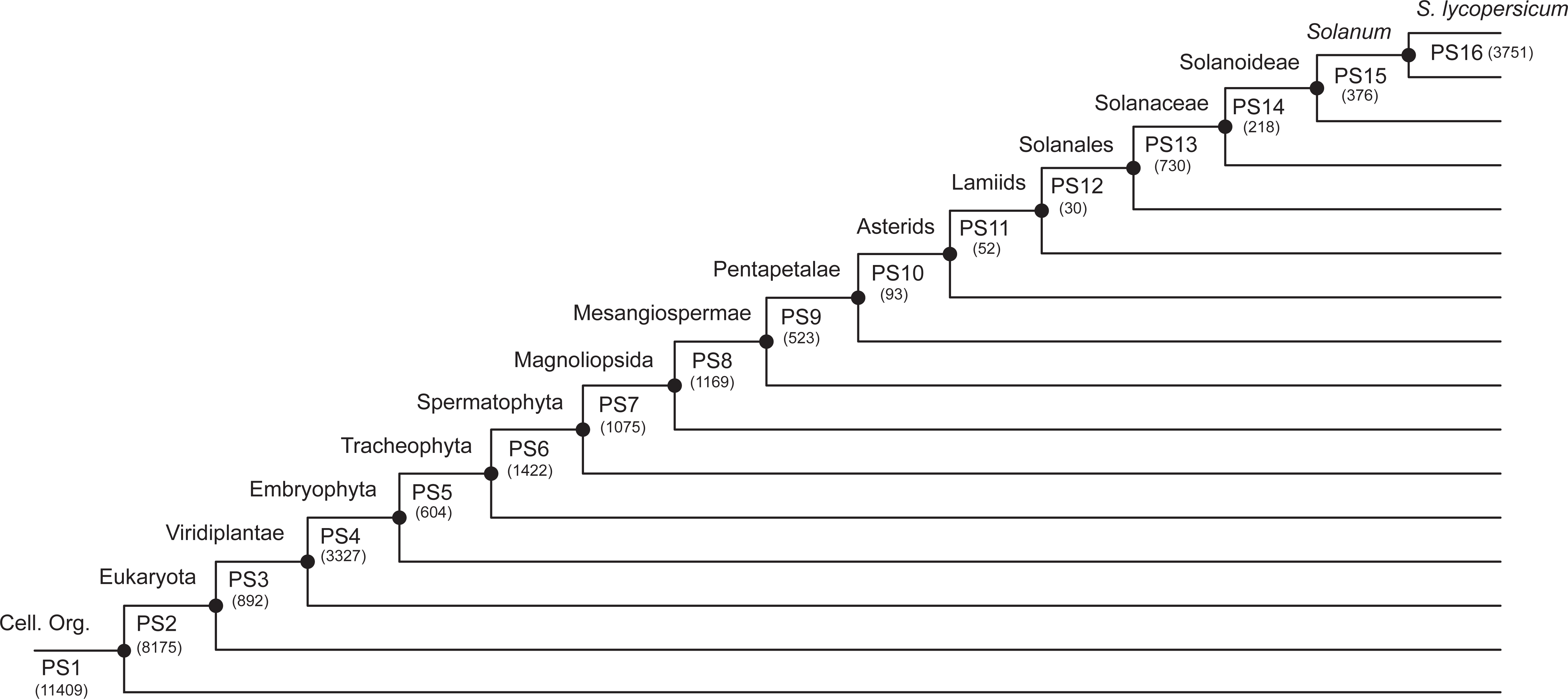
Phylostratigraphy of *S. lycopersicum* genes.

**Supplemental Fig. 10.**
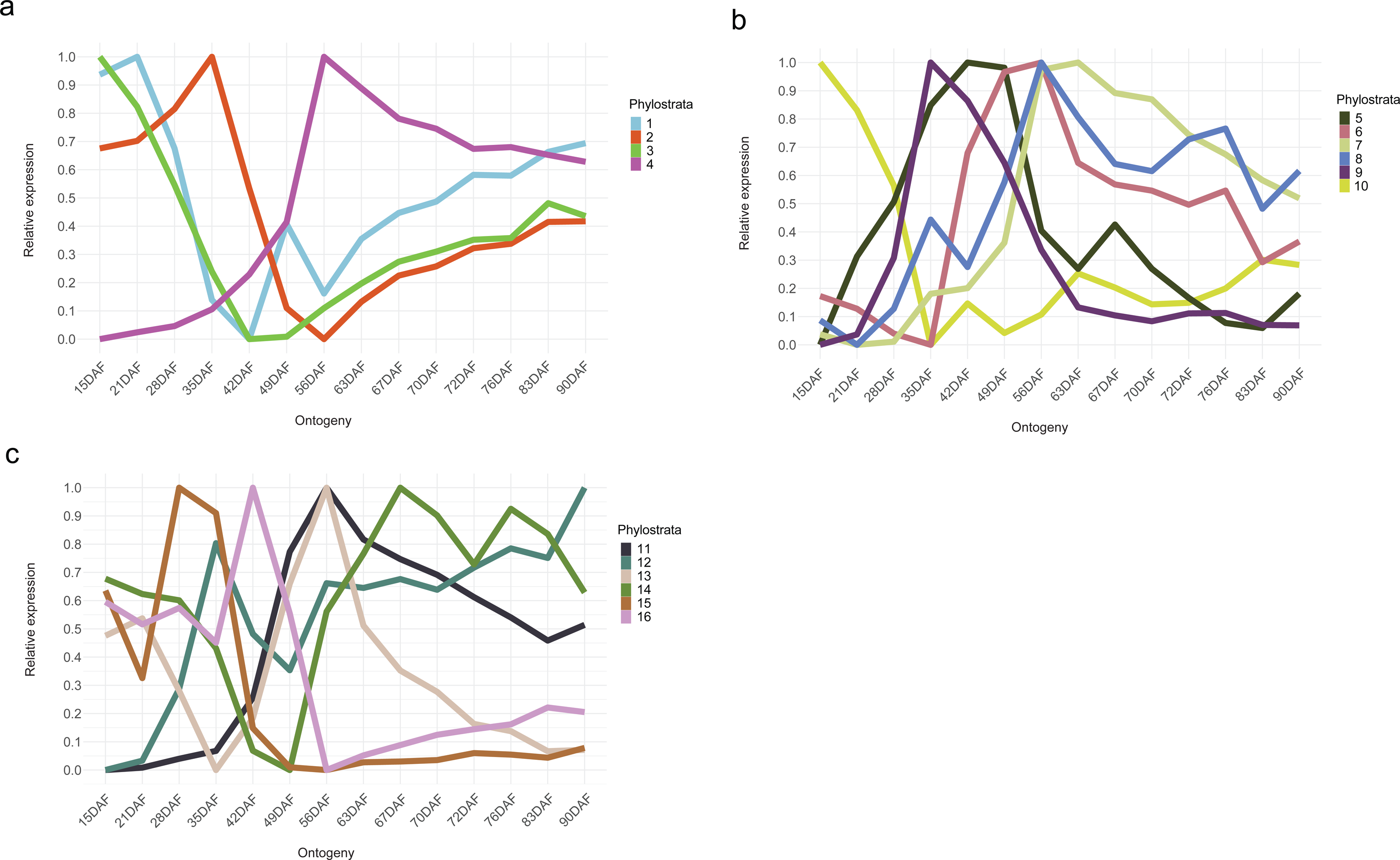
Relative expression of individual phylostrata during part of *S. lycopersicum* seed life cycle. **a**, PS1-PS4; **b**, PS5-PS10; and **c**, PS11-PS16. DAF indicates days after flowering.

**Supplemental Fig. 11.**
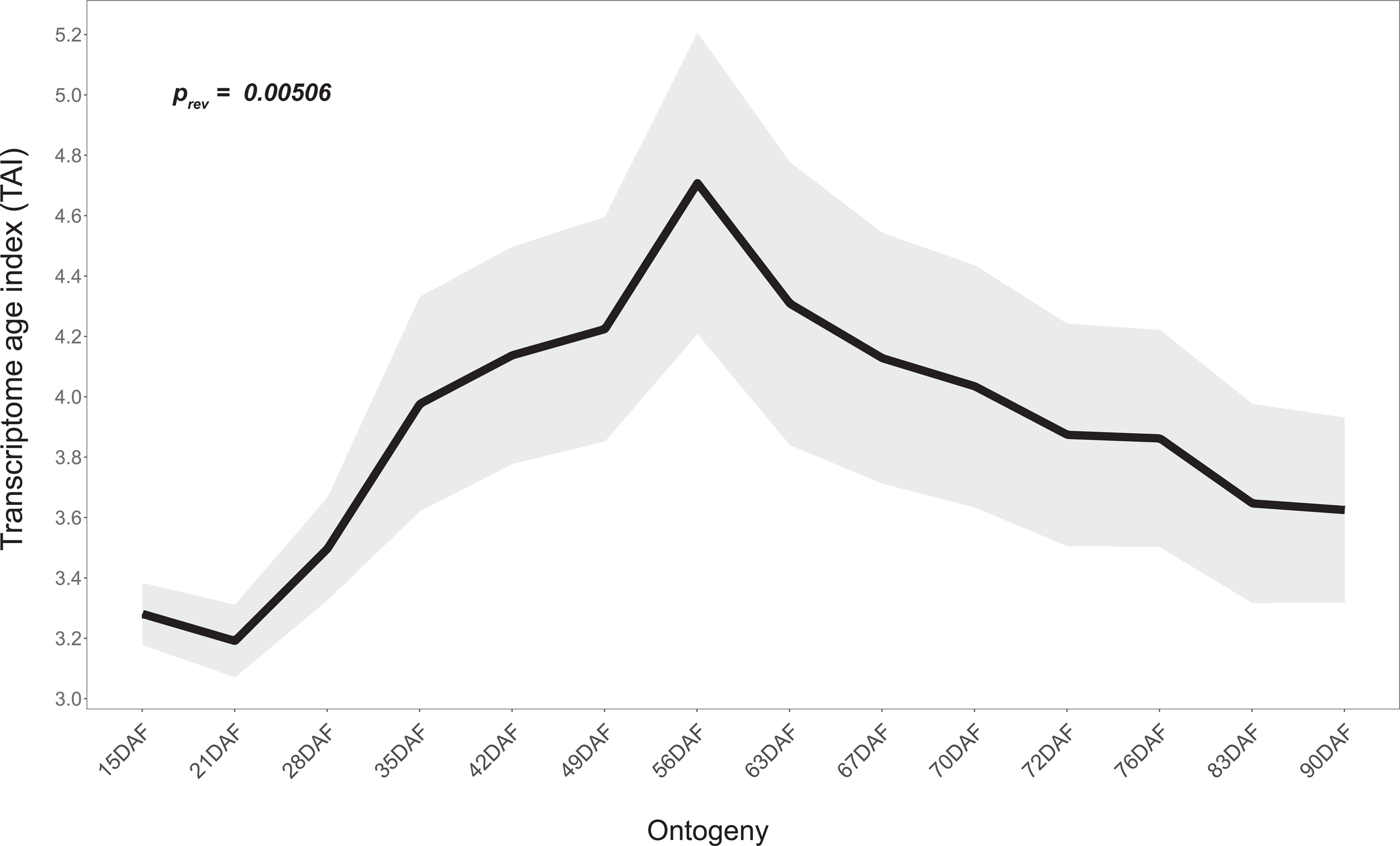
TAI profile during *S. lycopersicum* seed maturation excluding genes from PS16.

**Supplemental Fig. 12.**
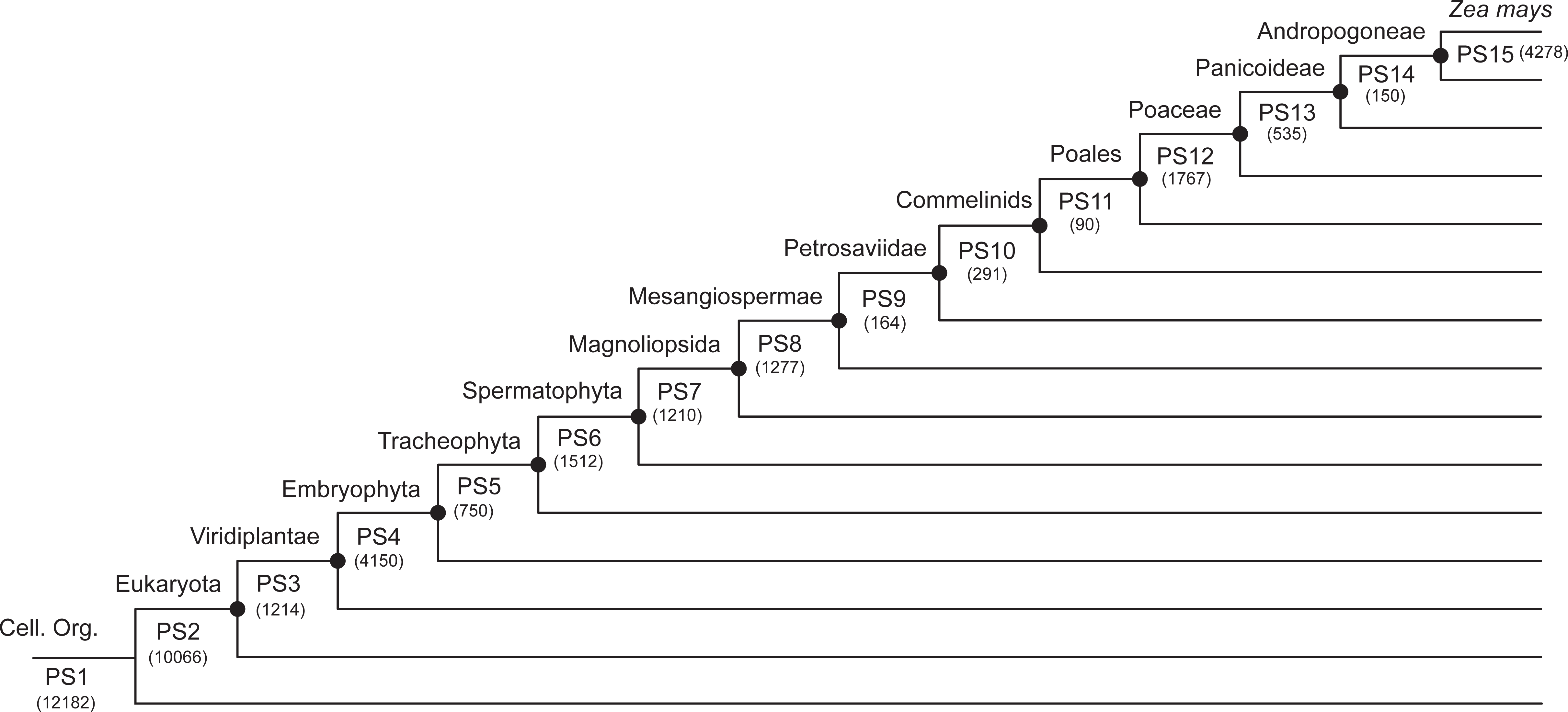
Phylostratigraphy of *Z. mays* genes.

**Supplemental Fig. 13.**
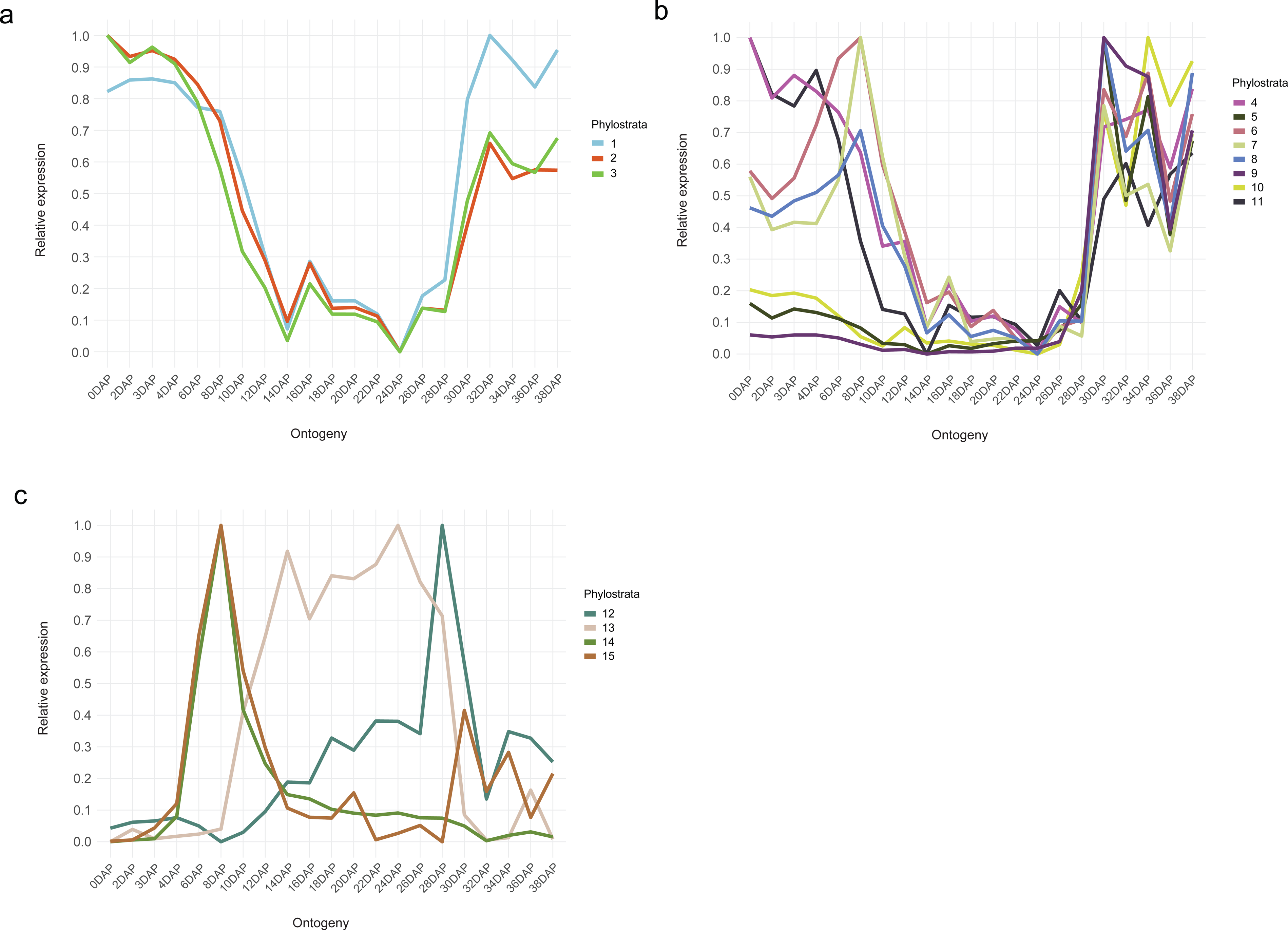
Relative expression of individual phylostrata during *Z. mays* seed life cycle. **a**, PS1-PS3; **b**, PS4-PS11; and **c**, PS12-PS15. Here, DAP refers to days after pollination.

**Supplemental Fig. 14.**
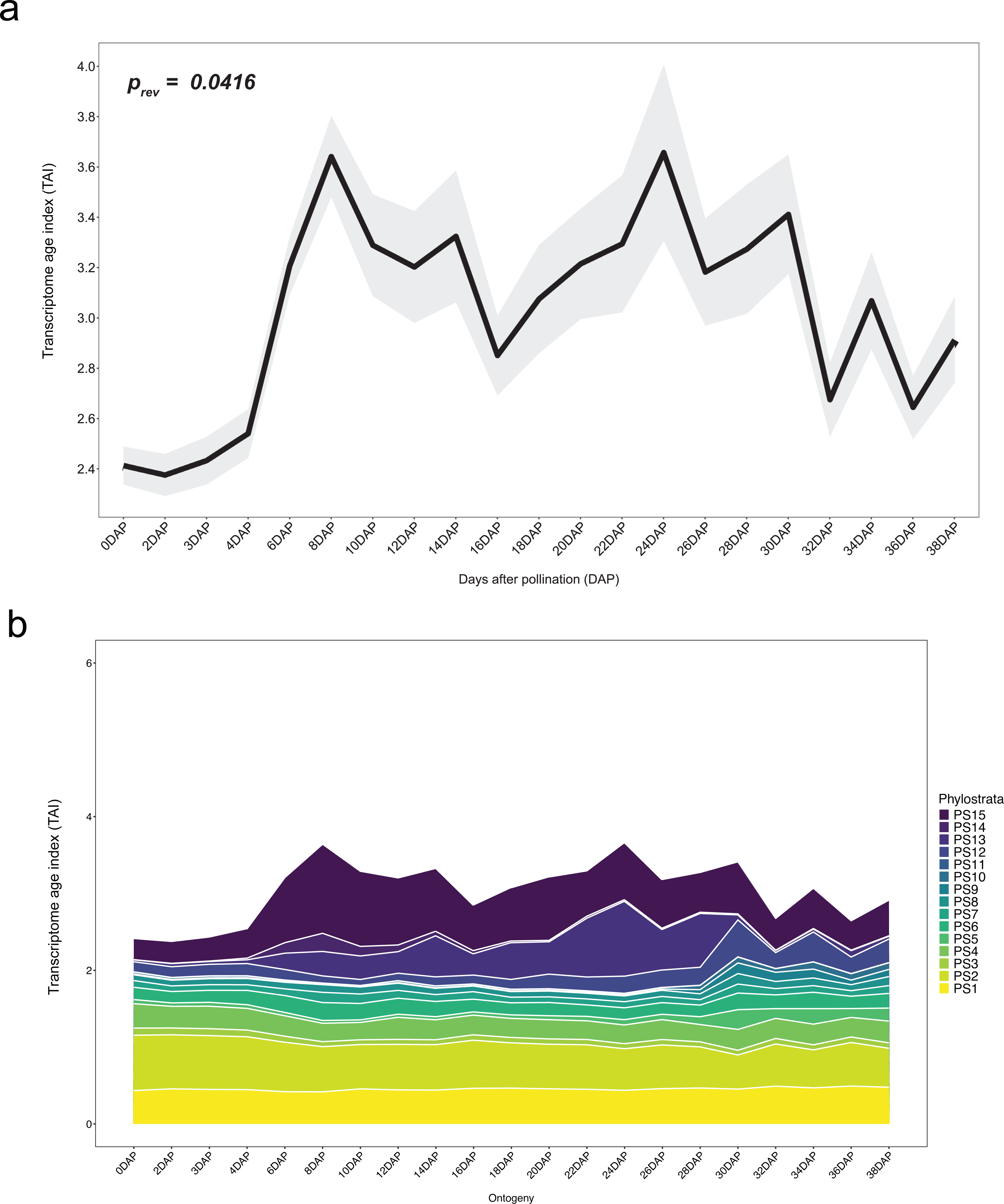
Phylotranscriptomics pattern of *Z. mays* without zein genes. **a**, TAI pattern over parts of the seed life cycle without zein genes. **b**, Individual phylostrata contribution to TAI profile without zein genes.

**Supplemental Fig. 15.**
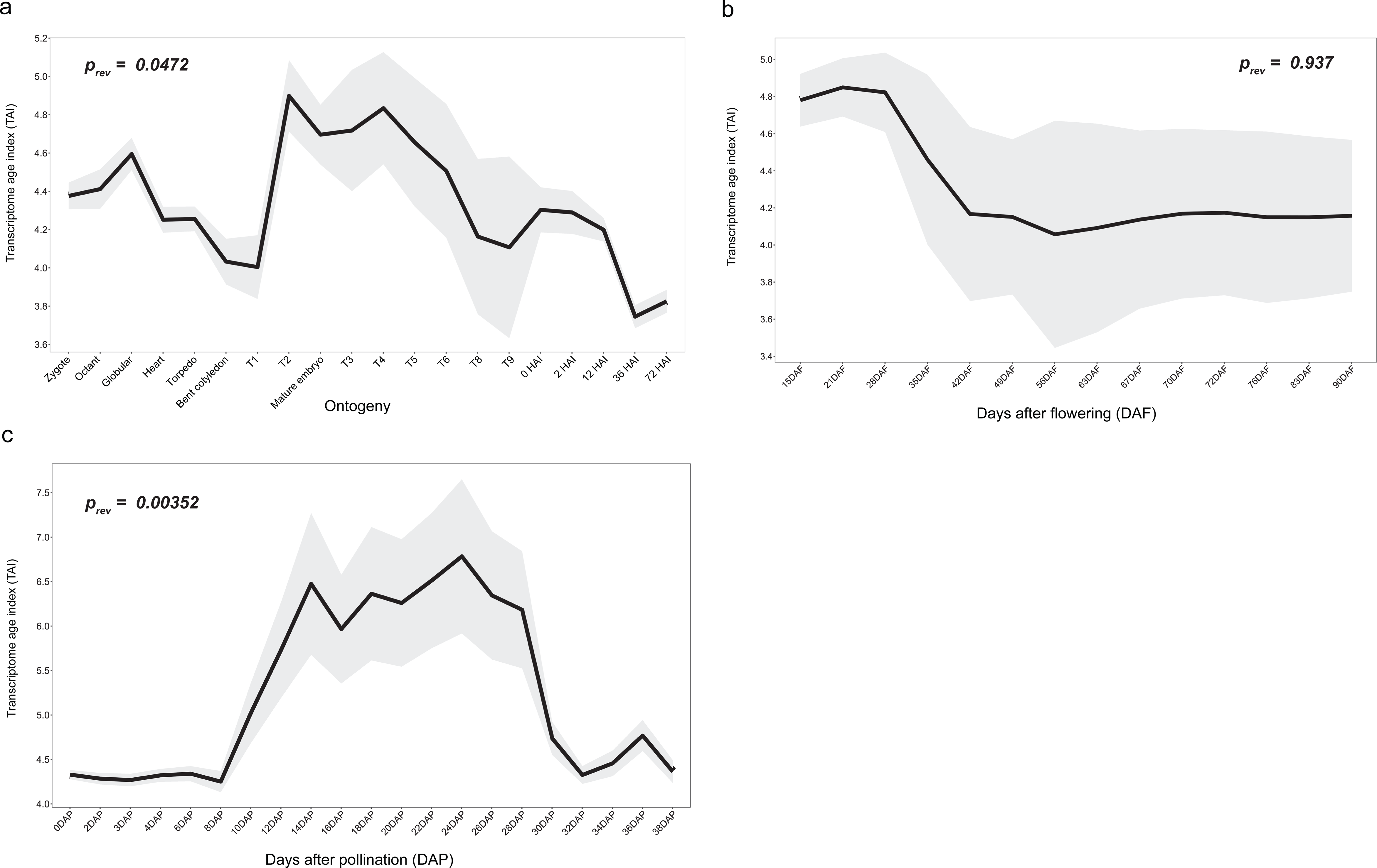
TDI pattern during parts of the seed life cycle in three angiosperm species - **a**, *B. napus*; **b**, *S. lycopersicum*; and **c**, *Z. mays*. The p-value was significant for a reverse hourglass test in *B. napus* and *Z. mays* but not in *S. lycopersicum*.

**Supplemental Fig. 16.**
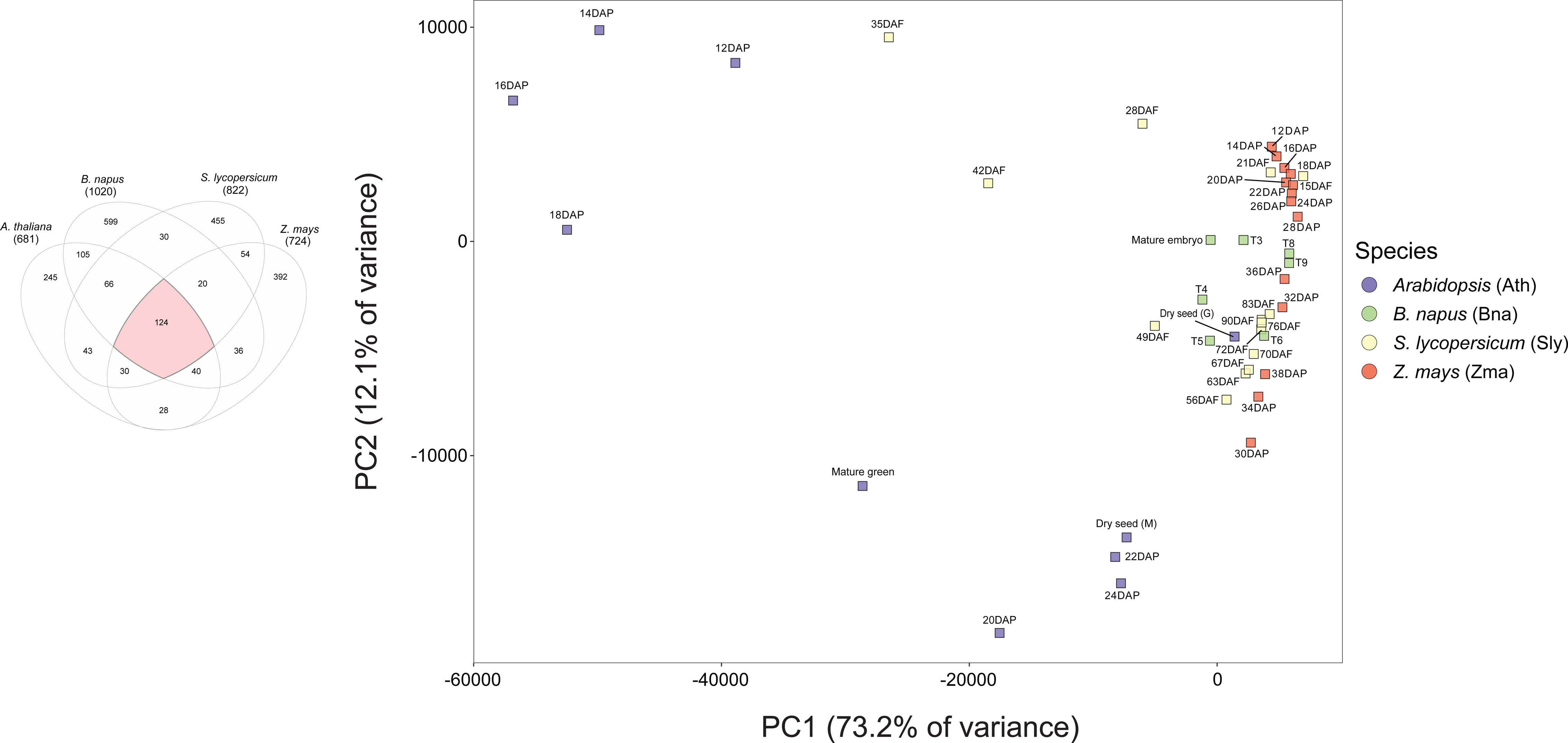
PCA based on the 124 shared top orthogroups (red area in Venn diagram). Time points from only the maturation phase from all four species were used for the PCA.

**Supplemental Fig. 17.**
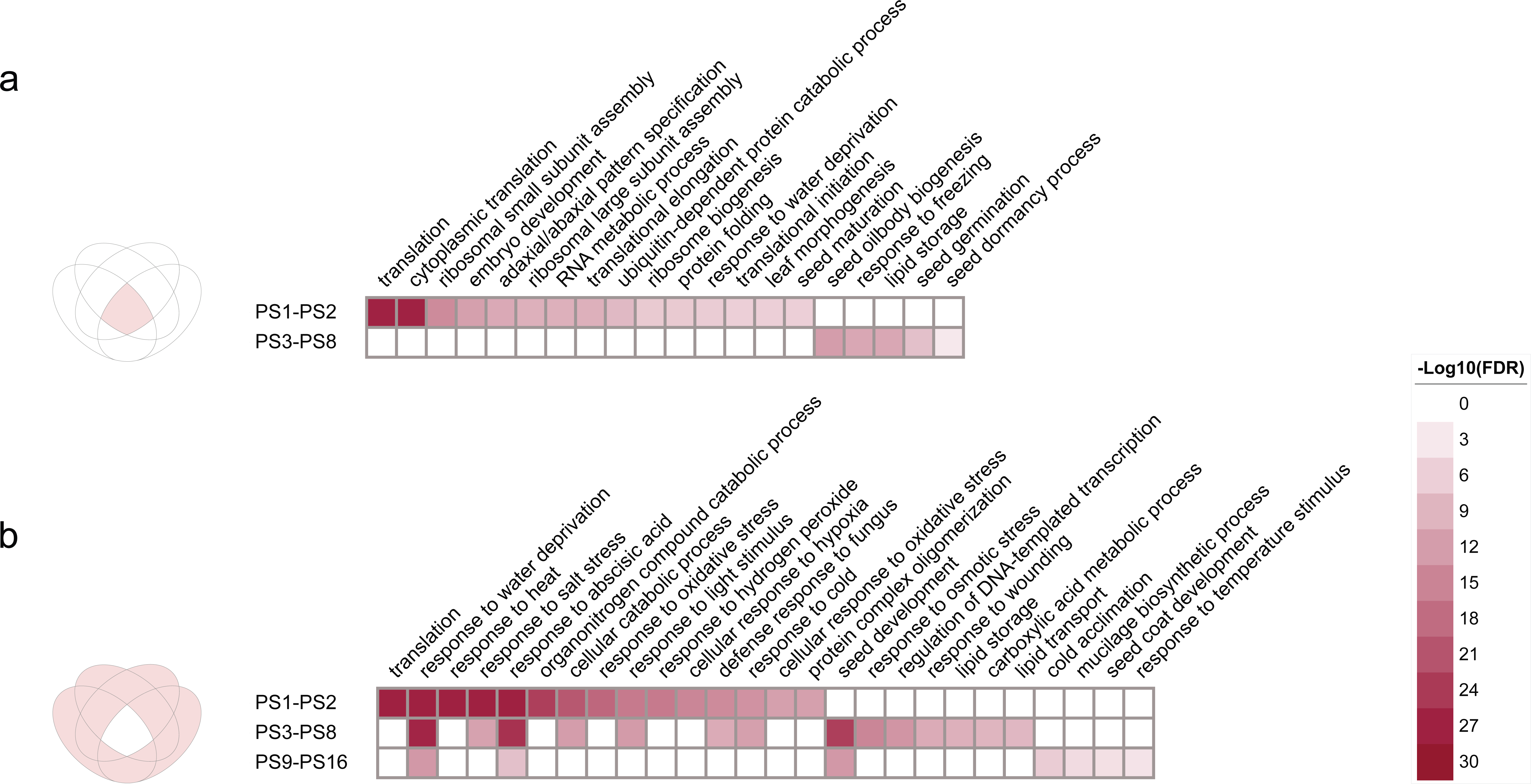
Top GO terms (sorted based on -log10(FDR)) enriched in *Arabidopsis* genes that belong to **a**, conserved top 5% orthogroups; and **b**, the rest of the orthogroups from the top 5%.

**Supplemental Fig. 18.**
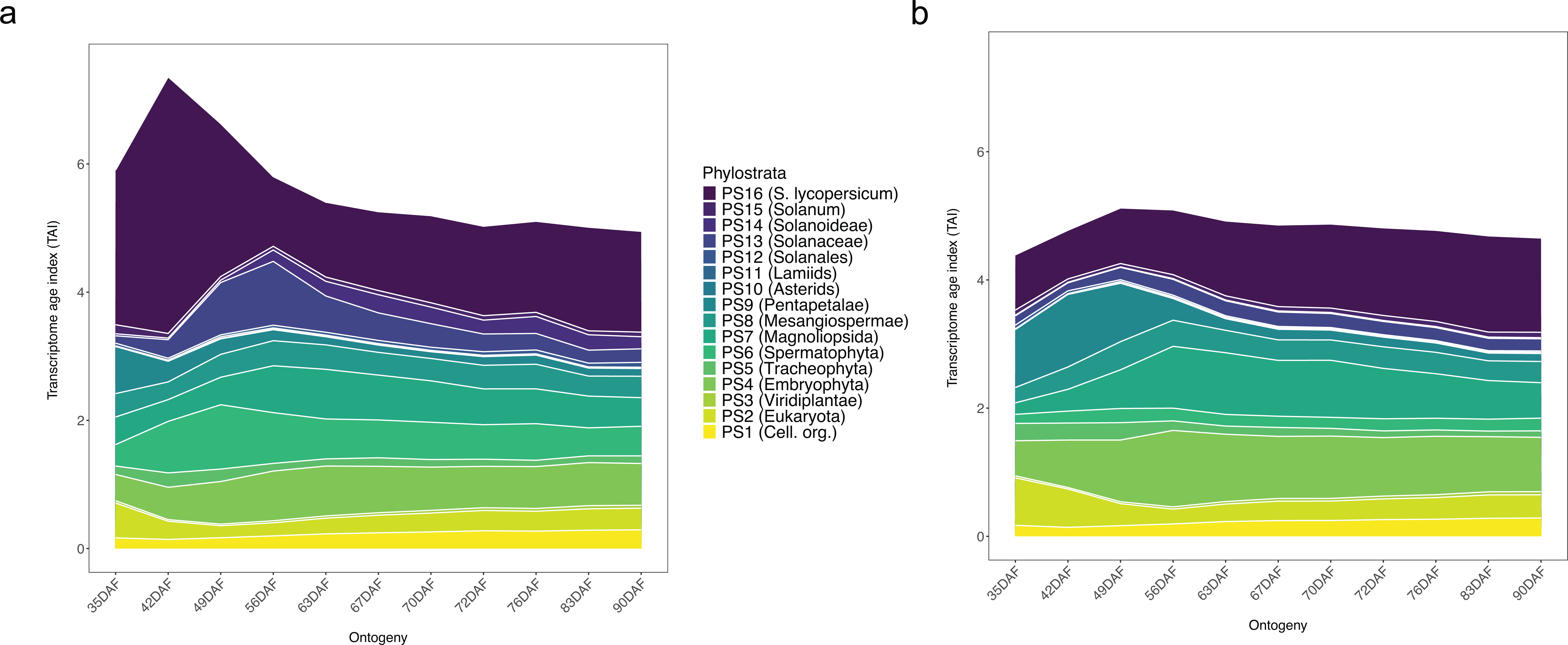
Tissue-specific individual phylostrata contribution to the overall observed TAI profile in *S. lycopersicum*. **a**, endosperm TAI; and **b**, embryo TAI.

**Supplemental Fig. 19.**
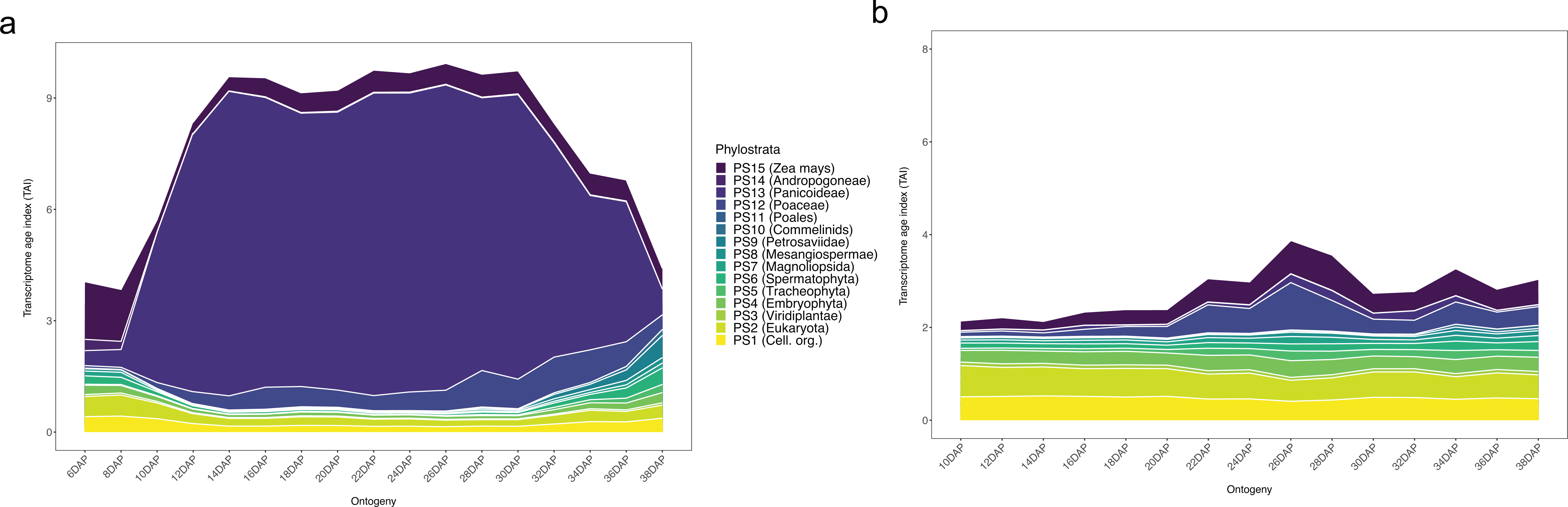
Tissue-specific individual phylostrata contribution to the overall observed TAI profile in *Z. mays*. **a**, endosperm TAI; and **b**, embryo TAI.

**Supplemental Fig. 20.**
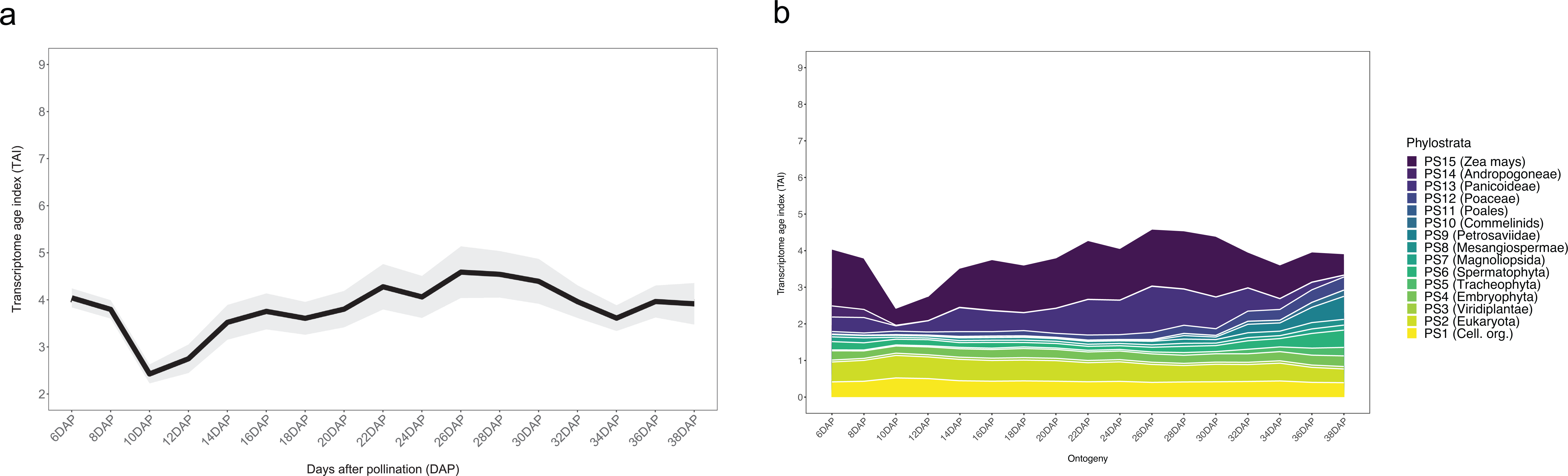
TAI profile of *Z. mays* endosperm tissue without zein genes.

**Supplemental Fig. 21.**
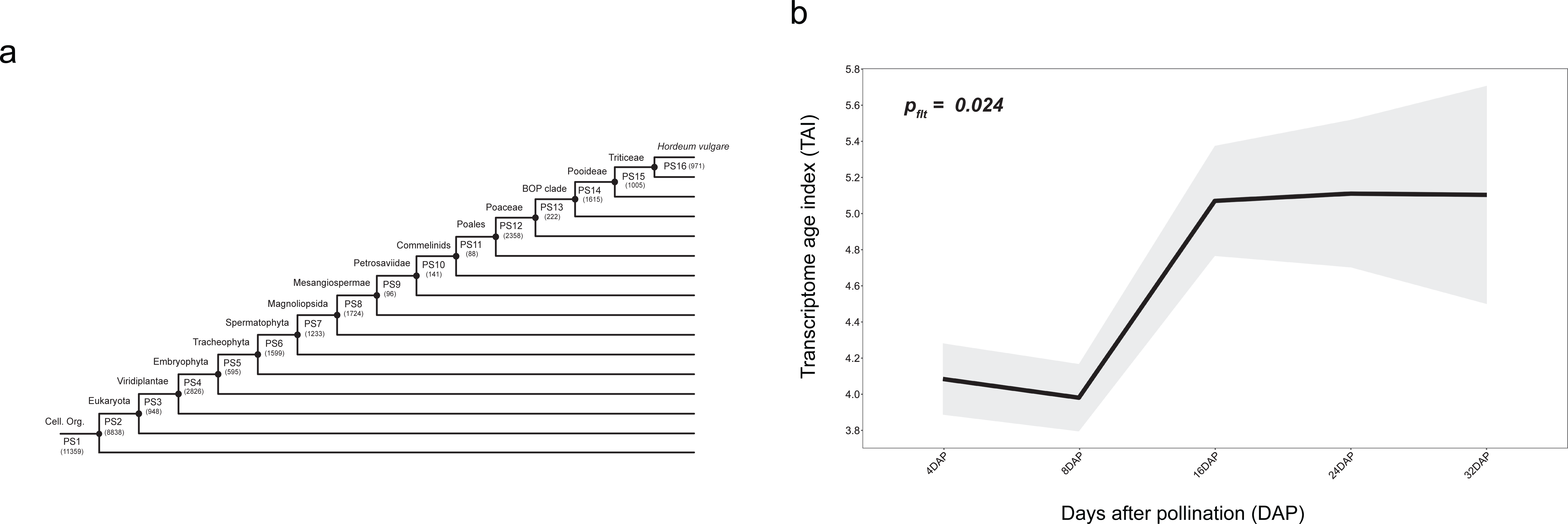
Phylotranscriptome of parts of *H. vulgare* seed life cycle. **a**, Phylostratigraphy of *H. vulgare* genes. **b**, TAI profile during part of *H. vulgare* seed life cycle.

**Supplemental Fig. 22.**
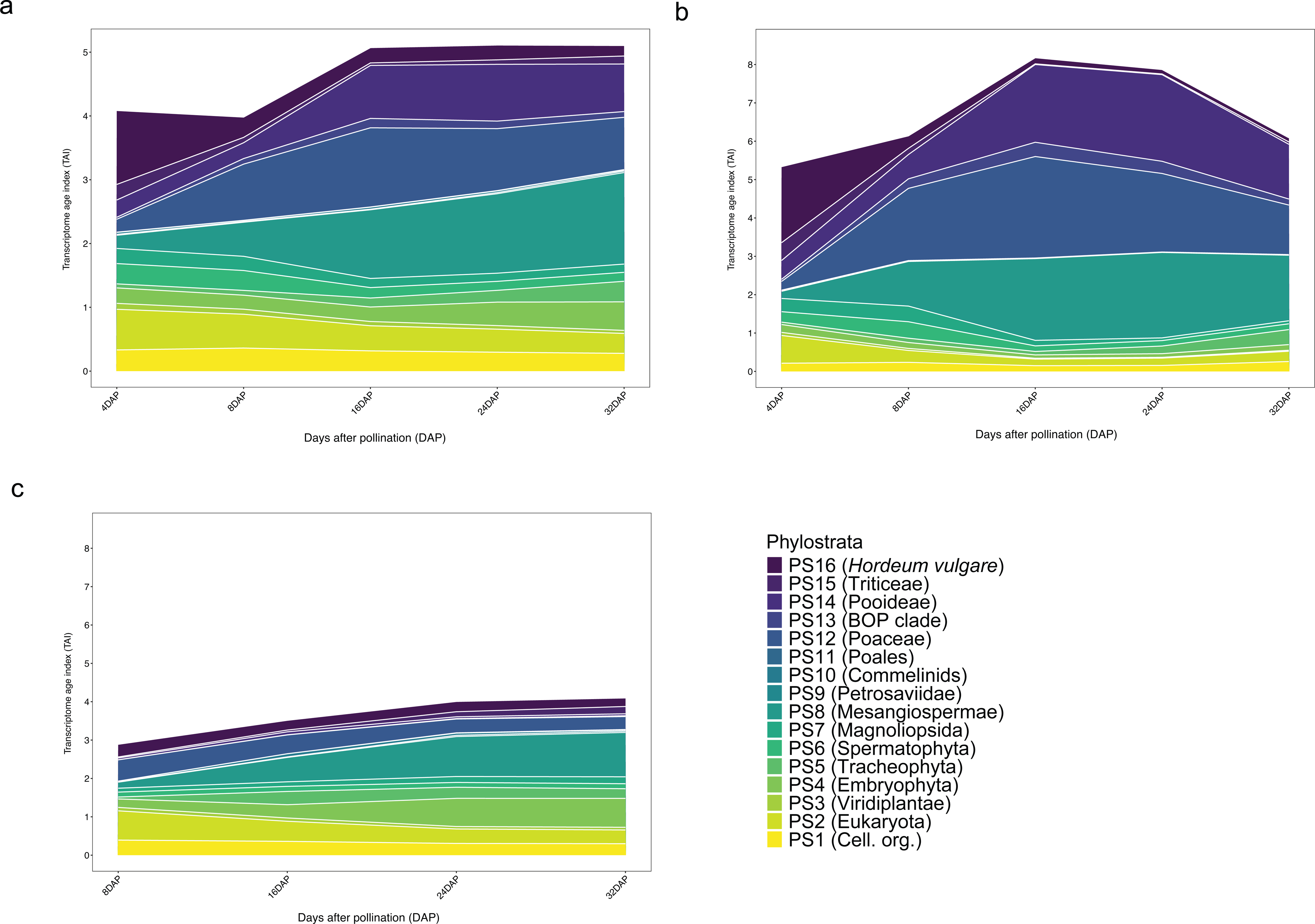
Contribution of individual phylostrata to the TAI profile of *H. vulgare* during part of the seed life cycle. **a**, TAI profile of the whole seed tissue. **b**, TAI profile of endosperm tissue. **c**, TAI profile of embryo tissue.

**Supplemental Fig. 23.**
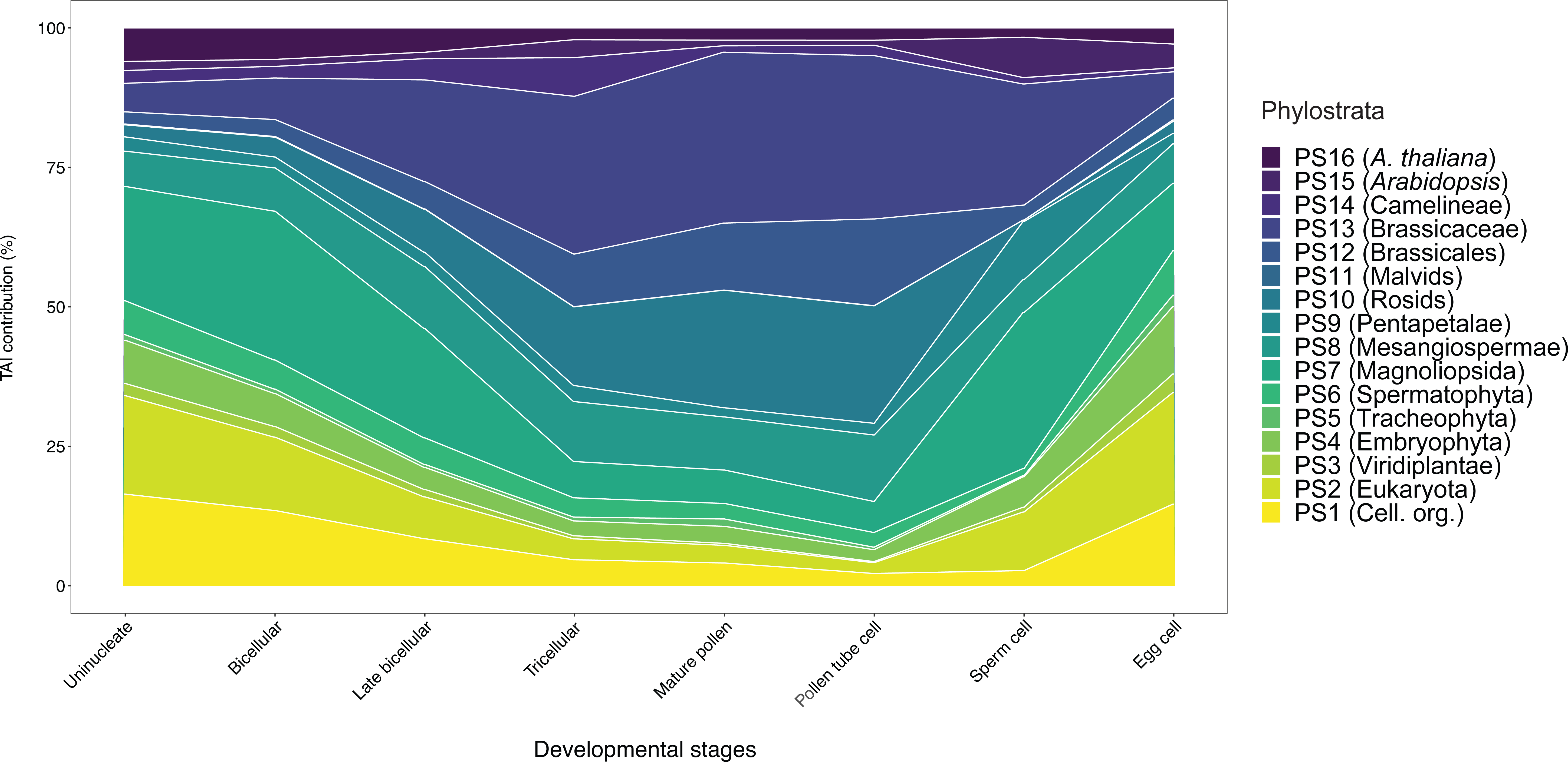
Percentage contribution of each phylostrata (PS1-PS16) to the overall TAI profile during *Arabidopsis* pollen development and egg cell stage.

**Supplemental Fig. 24.**
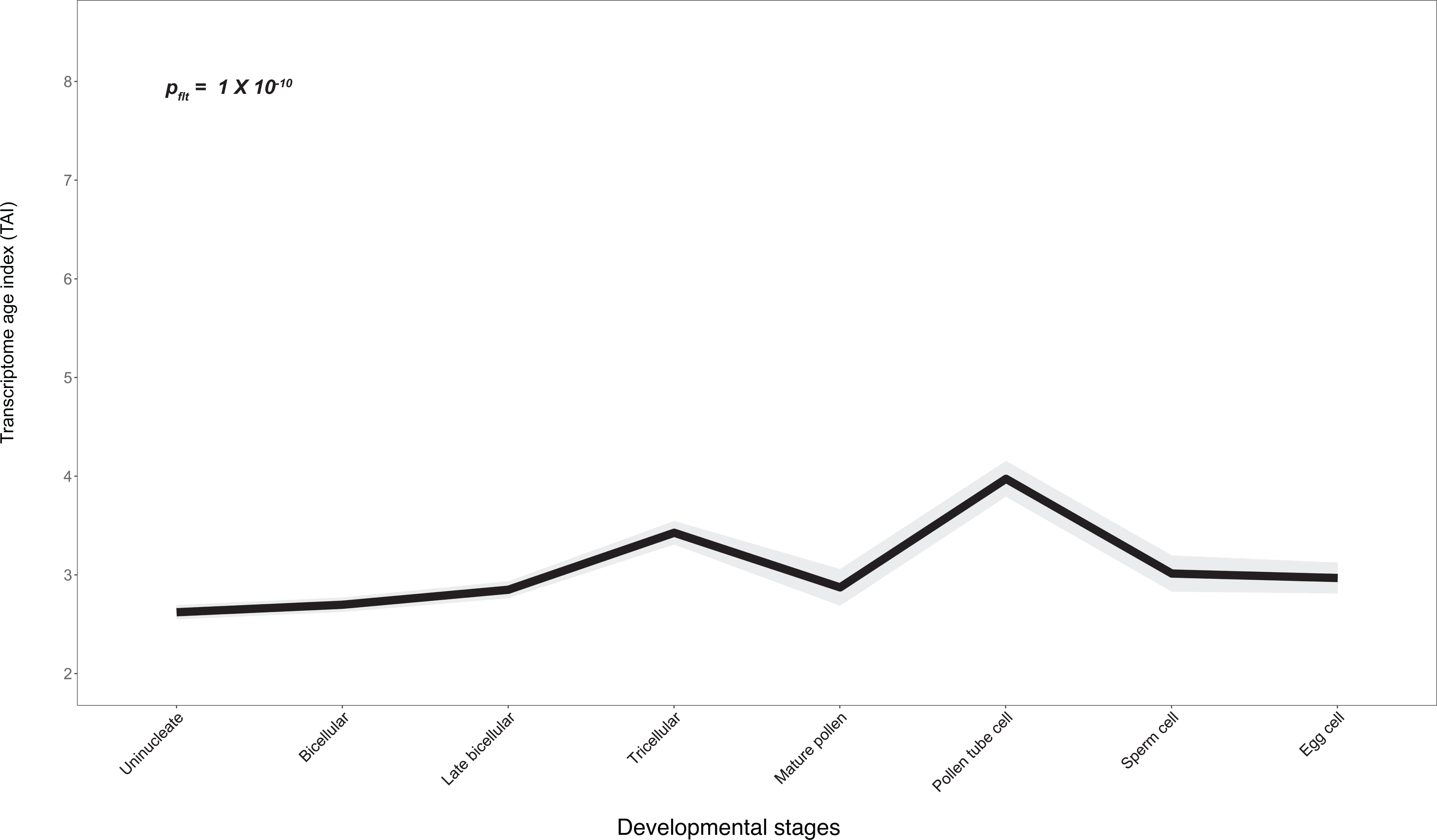
*Arabidopsis* pollen TAI profile without top 5% pollen expressed genes.

## Notes

### Competing Interest Statement

The authors have declared no competing interest.

